# Multi-scale Simulations of MUT-16 Scaffold Protein Phase Separation and Client Recognition

**DOI:** 10.1101/2024.04.13.589337

**Authors:** Kumar Gaurav, Virginia Busetto, Diego Javier Páez-Moscoso, Arya Changiarath, Sonya M. Hanson, Sebastian Falk, René F. Ketting, Lukas S. Stelzl

## Abstract

Phase separation of proteins plays a critical role in cellular organisation. How phase separated protein condensates underpin biological function and how condensates achieve specificity remain elusive. We investigated the phase separation of MUT-16, a scaffold protein in *Mutator foci*, and its role in recruiting the client protein MUT-8, a key component in RNA silencing in *C. elegans*. We employed a multiscale approach that combined coarse-grained (residue-level CALVADOS2 and near-atomistic Martini3) and atomistic simulations. Simulations across different resolutions provide a consistent perspective on how MUT-16 condensates recruit MUT-8, enabling the fine-tuning of chemical details while balancing the computational cost. Both coarse-grained models (CALVADOS2 and Martini3) predicted the relative phase separation propensities of MUT-16’s disordered regions, which we confirmed through *in vitro* experiments. Simulations also identified key sequence features and residues driving phase separation while revealing differences in residue interaction propensities between CALVADOS2 and Martini3. Furthermore, Martini3 and 350 µs atomistic simulations on Folding@Home of MUT-8’s N-terminal prion-like domain with MUT-16 M8BR cluster highlighted the importance of cation-*π* interactions between Tyr residues of MUT-8 and Arg residues of MUT-16 M8BR. Lys residues were observed to be more prone to interact in Martini3. Atomistic simulations revealed that the guanidinium group of Arg also engages in sp^2^-*π* interactions and hydrogen bonds with the backbone of Tyr, making Arg-Tyr interactions stronger than Lys-Tyr, where these additional favourable contacts are absent. In agreement with our simulations, *in vitro* co-expression pulldown experiments demonstrated a progressive loss of MUT-8 recruitment following the mutation of Arg in MUT-16 M8BR to Lys or Ala, confirming the critical role of Arg in this interaction. These findings advance our understanding of MUT-16 phase separation and subsequent MUT-8 recruitment, key processes in assembling *Mutator foci* that drive RNA silencing in *C. elegans*.

**Statement of Significance:** In cells proteins phase separate and form condensates. These protein condensates can play important role in bringing molecules together and facilitate biochemical processes. In this work, we used molecular dynamics simulations to understand how MUT-16 phase separates and forms the scaffold of the so-called *Mutator focus*. *Mutator foci* produce small RNA which help to regulates genes. As the scaffold of the *Mutator focus*, MUT-16 recruit multiple proteins which are important for the production of such small RNAs.

## Introduction

The formation of phase-separated condensates or membraneless organelles can organise cellular processes in time and space.^1^ The formation of such condensates is frequently underpinned by intrinsically disordered regions (IDRs) in multi-domain proteins or intrinsically disordered proteins (IDPs). IDRs and IDPs explore a myriad of different conformations rather than adopting a single well-defined 3D structure. IDRs and IDPs can engage in distributed, multivalent, and transient interactions that underpin the phase separation of protein and the formation of biomolecular condensates.^2,3^ Previous studies have suggested that protein phase separation is facilitated by different non-covalent interactions, including electrostatic, hydrophobic, sp^2^-*π*, and cation-*π* interactions between amino acids.^4–11^ Cation-pi interactions are formed between a positively charged cation of amino or guanidinium groups and *π* electron systems of aromatic groups within amino acid side chains.^12–15^ The interaction network facilitating the phase separation may also be at play in recruiting additional molecules to condensates.^16^ Frequently, a “scaffold protein” phase separates and recruits multiple “client” proteins, including enzymes that underpin the biological function of a condensate. Significant progress has been made in understanding interaction patterns that govern protein phase separation.^9,11,16^ However, a comprehensive understanding of how such interactions driving phase separation underpin molecular recognition on a molecular scale remains elusive.^17^

Membraneless organelles play a key role in RNA silencing, also called RNA interference (RNAi), an evolutionarily conserved gene regulation pathway in *C. elegans*. It is a primaeval defence mechanism to protect the genome from viruses and transposons and plays an essential role in gamete production, chromosome segregation, and development.^18,19^ In RNA silencing, small interfering RNAs (siRNAs) of 18-30 nucleotides are generated, recognising complementary messenger RNAs (mRNAs) to modulate their activity and stability.^19,20^ In *C. elegans*, RNA silencing requires amplification of primary siRNA to produce secondary siRNA facilitated by RNA-dependent RNA polymerases (RdRp)^21–24^ (Fig. 1A). This amplification occurs in perinuclear membrane-less organelles called *Mutator foci*.^25^ The loss of *Mutator foci* results in uncontrolled transposon proliferation and consequent sterility.^25,26^ *Mutator foci* are juxtaposed to P-granules responsible for RNA silencing and mRNA degradation (Fig. 1A). The proximity of another membraneless organelle suggests the need for a mechanism that ensures specific molecular recognition.

**Figure 1:**
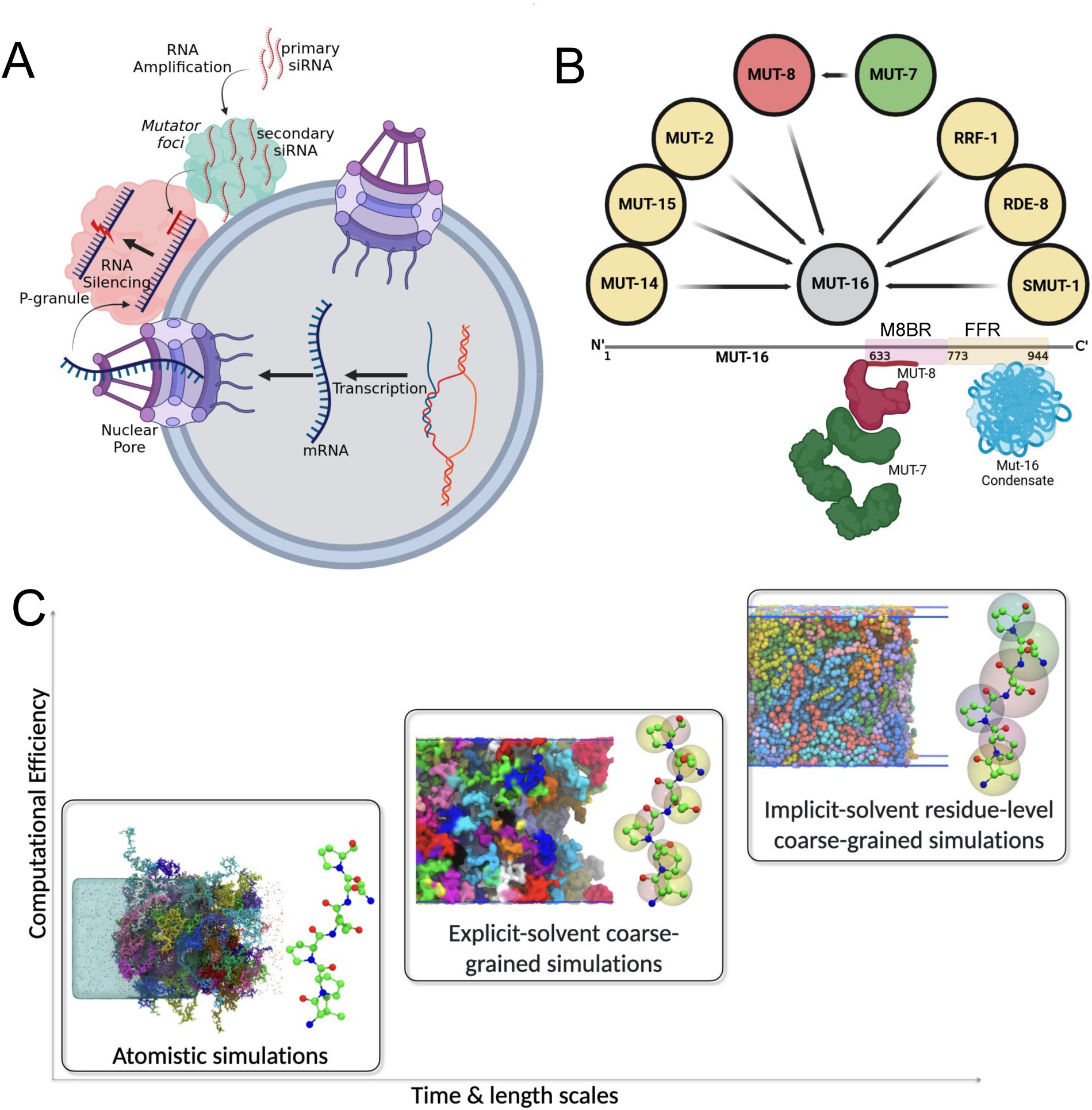
**A.**RNA silencing in *C. elegans*, involving siRNA amplification within membraneless *Mutator foci*. Generated siRNAs move to P-granules, directing the RNA-induced silencing complex (RISC) to target specific mRNAs for degradation. **B.** The scaffold protein MUT-16 (grey) assembles Mutator foci, recruiting essential siRNA amplification proteins (yellow), including MUT-8 (red), which subsequently recruits the exoribonuclease MUT-7 (green). Regions of MUT-16 responsible for phase separation (FFR, yellow) and MUT-8 binding (M8BR, pink) are shown. The MUT-8 N-terminal (red) interacts with MUT-16 M8BR, while its C-terminal binds MUT-7. MUT-16 FFR (blue) is shown forming phase-separated condensates. **C.** Multi-scale simulation framework to investigate phase behavior and molecular recognition, combining atomistic, near-atomic (Martini3), and residue-level (CALVADOS2) simulations.

The MUT-16 protein, which features extended disordered regions (Fig. S1A), is the scaffold for the *Mutator foci*. MUT-16 foci dissolve in living *C. elegans* when the temperature is elevated and reappears on lowering the temperature. The temperature response *in vivo* is consistent with an Upper critical solution (UCST) phase behaviour, but further characterization of MUT-16 is required to understand how they form, whether this is really protein phase separation and what interactions are driving MUT-16 foci formation. Notably, the C-terminal region (773-1054 aa) is crucial for foci formation *in vivo*.^25^ Different regions of MUT-16 recruit proteins required for various steps of siRNA amplification such as the RNA-dependent RNA polymerase RRF-1, the nucleotidyltransferase MUT-2, the DEAD-box RNA helicase MUT-14, the 3’-5’ exoribonuclease MUT-7, and MUT-8 (also known as RDE-2)(Fig. 1B).^25,27^ The exoribonuclease MUT-7, a critical enzyme in transposon repression within *C. elegans*,^26^ is recruited to the *Mutator foci* via its interaction with MUT-8,^28^ which serves as a linker between MUT-7 and the scaffolding protein MUT-16. MUT-16 features a designated MUT-8 binding region (M8BR, residues 633–772) responsible for recruiting MUT-8 (Fig. 1B).^29^ Interestingly, a prion-like domain is present in the MUT-8 N-terminus (Fig. S1B). Prion-like domains (PLDs) are often associated with protein phase separation, and the MUT-8 PLD could potentially facilitate its co-condensation with MUT-16. However, the potential interaction between MUT-16 and the MUT-8 N-terminal PLD and the molecular mechanisms facilitating this interaction remains unknown.

Sequence drivers of phase separation and specific molecular recognition between disordered proteins can be revealed by simulations at different scales from highly coarse-grained to atomistic simulations.^30–33^ However, the extent of agreement between these simulation approaches remains uncertain. Residue-level implicit solvent simulations with single-bead-per-amino-acid models, such as Mpipi^34^ and CALVADOS2,^35,36^ are computationally efficient and proficient at capturing the relative phase separation propensities of disordered proteins. These methods have enabled proteome-scale investigations of intrinsically disordered regions (IDRs).^37,38^ Increasing the resolution (Fig. 1C), for instance, to a near-atomic resolution explicit solvent Martini3 simulations, ^39^ has the potential to capture interactions between different chemical moieties better. The Martini3 model enables simulations of many different types of molecules but has not been developed to model protein phase separation.^40–43^ Nonetheless, with a small adjustment of the relative strengths of protein-water interactions, Martini3 models accurately capture trends for conformational ensembles in IDRs in multi-domain proteins and IDPs.^42–44^ Consequently, it may be possible to use Martini3 to gain additional insights into the protein condensates at the expense of more extensive computational requirements. Increasing the computational demands even further, atomistic molecular dynamics simulations provide detailed insights into specific molecular interactions of individual amino acids within condensates.^11,31–33,45,46^

Although multi-scale simulation methods have become increasingly established, it is less clear to what extent, e.g., coarse-grained simulations at one-bear-per amino acid resolution CALVADOS2 and near-atomic resolution Martini3 simulations would give consistent results for a given protein. Differences in their parameterisations might lead to differences for specific proteins in terms of which regions of the proteins are predicted to drive interactions and how well coarse-grained simulations match experimental trends. Ideally, we would expect that sequence features identified with residue-level implicit solvent simulations would also be seen in more highly-resolved near atomistic resolution simulations. Overall, coarse-grained simulations would capture long timescales that are computationally expensive in atomistic simulations, providing insights into sequence features and interaction patterns that align with those identified in atomistic models. In practice, atomistic simulations offer fine-grained detail, enabling identifying interactions that may be over- or underestimated in coarse-grained models, thus adding accuracy to the interpretation of molecular recognition and phase behaviour across scales. It is unclear whether simulations at different scales^47^ provide consistent predictions. Increasing the resolution from coarse-grained models with implicit solvents to explicit solvent models adds more chemical detail. However, it is unclear if this change affects essential features of the system.

In this study, we elucidate how the MUT-16 phase separates and recruits MUT-8 N-terminal PLD through a comprehensive multi-scale simulation approach (Fig. 1C), complemented by *in vitro* experiments that report on phase behaviour and protein-protein interactions. We systematically elevate the resolution of our simulation models, progressively transitioning from amino acid bead-string representations to more intricate coarse-grained models that approach near-atomic resolution, eventually culminating in fully atomistic molecular dynamics simulations. Our coarse-grained simulations, using the CALVADOS2 and Martini3 models alongside *in vitro* experiments, emphasise the critical role of the MUT-16 fociforming region (FFR, 773-945 aa) in phase separation. The simulations further identified key residues, including Tyr, Arg, Phe, Pro, Gln, and Asn, as major contributors to this process. Through Martini3 and 350 µs atomistic simulations of the MUT-16 M8BR complex with MUT-8 fragments, we demonstrate that Arg-Tyr pairs, synergistically stabilized by cation-*π*, sp^2^-*π*, and hydrogen bonds, play a key role in recruiting the MUT-8 N-terminal PLD to the MUT-16 condensate. This finding was validated through *in vitro* co-expression pulldown experiments using MUT-8 and wild-type MUT-16 M8BR, as well as Arg mutants (R *>* K and R *>* A) of MUT-16 M8BR. The Arg mutants (R *>* K and R *>* A) of MUT-16 M8BR exhibited a progressive reduction in MUT-8 binding compared to wt MUT-16 M8BR, with the R *>* K mutant showing a moderate loss and the R *>* A mutant displaying a more pronounced loss. These results confirm the crucial role of MUT-16 M8BR Arg in recruiting MUT-8.

## Methods

We performed molecular dynamics simulations in three resolutions (Fig. 1C): implicit solvent residue-level coarse-grained simulations (CALVADOS2),^35,36^ near-atomic explicit-solvent coarse-grained simulations (Martini3),^39,40^ and atomistic simulations. Combining the simulations at three resolutions enables us to identify residues engaging in key interactions that favour MUT-16 condensate formation and MUT-8 recruitment to MUT-16 condensates and then zoom in on these interactions in atomistic molecular dynamics. Further, we performed *in vitro* experiments to investigate and validate the MUT-16 phase separation and the recruitment of the MUT-8 N-terminal domain.

### Residue-level coarse-grained simulations

Residue-level coarse-grained simulations were executed using the CALVADOS2 model with GPUs using the HOOMD-blue software package (v. 2.9.6)^48^ extended with azplugins (v. 0.10.2). The CALVADOS2 model is an improvement of the previously established HPS model,^30^ proposed by Tesei et al.^35,36^ Within the framework of the CALVADOS2 model, each amino acid is treated as a single bead suspended in an implicit solvent environment. This model enables the investigation of sequence-specific interactions among biomolecules and effectively circumvents the temporal limitations associated with higher-resolution models. The potential energy function employed for the simulations encompasses bonded and non-bonded components, incorporating electrostatic and short-range pairwise interactions. A harmonic potential describes the bonded interactions,

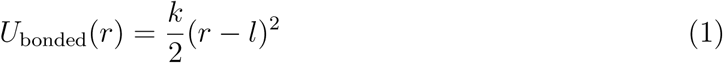

where the spring constant k = 8368 kJ mol*^−^*^1^nm*^−^*^2^ and equilibrium bond length l = 0.38 nm. The non-bonded interaction between the monomer beads is described by Ashbaugh-Hatch potential,^49^

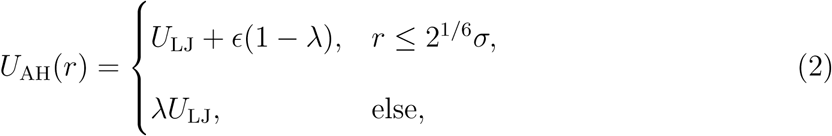

where *E* = 0.8368 kJ mol*^−^*^1^ and *U*_LJ_ is the Lennard-Jones potential:

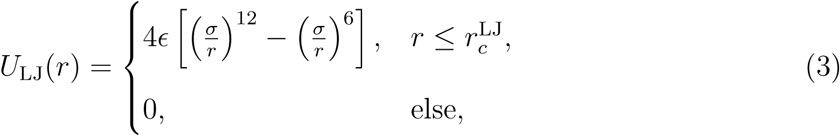

where 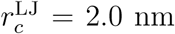.^36^*σ* and *λ* are determined by computing the arithmetic average of amino acid specific parameters denoting size and hydrophobicity, respectively. The residue-specific *λ* were previously optimised by Tesei et al., with a Bayesian method that uses a comprehensive experimental data set. ^35,36^ The non-bonded interaction also includes an electrostatic component modeled via Debye-Hückel potential:

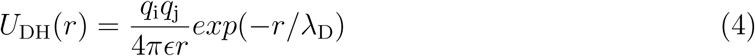

where *q*_i_ and *q*_j_ are charges. The Debye screening length (*λ*_D_) and the dielectric constant (*E*) are set to 1 nm and 80, respectively to reproduce the physiological conditions. The electrostatic potential is truncated at *r_c_* = 4.0 nm.

The simulations were performed with 100 protein chains in the slab geometry of size 20 × 20 × 280nm under periodic boundary conditions (See Table S1).^30^ The simulations were initialised by placing the protein chains randomly in the slab. The Langevin thermostat regulated the temperature. Further, the equations of motions were integrated with a timestep (Δ*t*) of 10 fs. The simulations were typically performed for 10 µs (Table S1).

### Phase diagram from residue-level coarse-grained simulations

The residue-level coarse-grained (CALVADOS2)^35^ simulations were performed at different temperatures (260 K, 265 K, 270 K, 275 K, 280 K, 285 K, 290 K, 291 K, 292 K, 293 K, 295 K, 300 K) for 10 µs each (Table S1). To determine the densities of dense and dilute phases, we followed the approach by Tesei et al.^35^ The protein concentration of dilute (*ρ*_l_) and dense (*ρ*_h_) phases are calculated along the z-axis. The critical temperature *T_c_*was obtained by fitting the densities obtained from the simulation to:

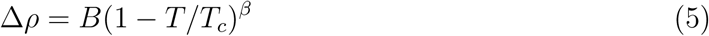

where Δ*ρ* = *ρ*_h_ − *ρ*_l_, B is critical amplitude and *β* = 0.325 is critical exponent. Further we obtained the critical density *ρ*_c_ by fitting the simulations data to:

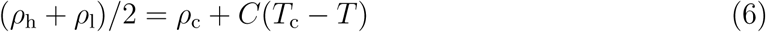

where *C* is a positive fitting parameter.^30^

### Near-Atomic Coarse-Grained Simulations with Explicit Solvent Using the Martini3 Model

The near-atomic coarse-grained (Martini3) simulations were executed using GROMACS,^50^ employing either the latest version of the Martini force field (Martini3),^39,40^ or an improved variant of Martini3 featuring rescaled protein-water interactions.^42^ In the Martini3 force-field framework, multiple atoms are grouped into a single particle, often representing four to five heavy atoms with a single coarse-grained bead.^39^ Water and ions are represented explicitly. The initial structure of the protein IDR chains was obtained by taking in desired regions from the AlphaFold^51^ structures of MUT-16 (AF-O62011-F1) and MUT-8 (AF-Q58AA4-F1). The protein chains were coarse-grained using the martinize2 python script.^52^ The coarse-grained protein chains were inserted in a cubical box or a slab geometry and further solvated with the Insane.py Python script.^53^ Furthermore, 0.15 M NaCl was added to the system on top of balancing the charge. Simulations were performed with periodic boundary conditions. The simulation box was subjected to minimisation using the steepest descent algorithm. Firstly, the water and salt were equilibrated in the NVT ensemble by applying the position restraint on the protein. Secondly, the system was further equilibrated by removing the position restraint on the protein in the NVT ensemble. Thirdly, the system was subjected to equilibration in an NPT ensemble for 700000 steps with a 20 fs time-step. Finally, production simulations were performed in the NPT ensemble. The final temperature was maintained at 300 K using the Bussi-Donadio-Parrinello velocity-rescaling thermostat,^54^ and the pressure was maintained at 1 bar using Parrinello-Rehman barostat.^55^ All the simulations were performed for 20 µs with a 20 fs time step. Last 10 µs of the trajectory was used for the analysis.

### Density calculations from Martini3 simulations

Radial density profiles were calculated to investigate the spatial distribution of the MUT-16 M8BR+FFR, MUT-16 FFR and MUT-16 M8BR chains relative to the center-of-mass (COM) of the system. The methodology was adapted from Benayad et al.^44^ Each bin had a width of 10 Å.

### Contact-map analysis

Two-dimensional (2-D) inter-chain contact maps were generated by calculating contact probabilities between residues. First, distances between all residue pairs across protein chains were measured. These distances were then converted into a contact matrix, with values set to 1 for distances below the defined cutoff and 0 for those above it. The pair cutoff was set to 2^1/6^*σ_ij_*,^30^ 6 Å, and 4.5 Å for CALVADOS2, Martini3, and atomistic simulations, respectively. The resulting contact matrix was averaged over all protein chain pairs and further averaged across simulation frames to produce the final contact map. For atomistic and Martini3 simulations, the contact analysis was based on the procedure described in https://github.com/dwhswenson/contact_map and for CALVDAOS simulations MD-Analysis was used.^56,57^ One-dimensional contact maps were computed by averaging the values of the final contact matrix along the respective axis. A region-wise contact map was acquired by averaging the values within the specific region of interest from the final contact matrix. On the other hand, the residue-wise contact map was generated by summing the values of each residue pair (e.g., Arg-Tyr, Phe-Met) obtained from the final contact matrix and normalising them with the corresponding number of residue pairs.

### Atomistic molecular dynamics simulations

The atomistic simulations were performed for the MUT-16/MUT-8 interaction system obtained from the last frame of the Martini3 simulation trajectory. The system was backmapped using backmap.py python script.^58^ For computational efficiency, we reduced the slab dimensions from 15 nm× 15 nm × 60 nm to 15 nm×15 nm×60 nm by selectively removing water molecules from the z-direction. The atomistic simulation system features 884,932 atoms. Simulation was performed using the Amber99sb-star-ildn-q force field^59–62^ with TIP4P-D water^63^ in GROMACS.^50^ The simulation box contained 100 MUT-16 M8BR and 10 MUT-8 N-terminal chains. After backmapping, the atomistic system was minimised using the steepest-descent algorithm to remove clashes. Further, the system was equilibrated in three steps similar to Martini3 simulations. Temperature (300 K) and pressure (1 bar) were maintained using the Bussi-Donadio-Parrinello velocity rescaling thermostat^54^ and Parrinello-Rahman barostat.^55^ The production run was performed for 1 µs in the NPT ensemble in in-house supercomputer Mogon-II and Further, 100 parallel simulations totalling 350 µs in Folding@home^64^ were launched from 100 different conformations taken from the 1 µs trajectory. For each conformation, 100 simulations with different random velocities were launched.

### Cation-*π* and sp^2^-*π* interactions

Cation-*π* interactions between Arg/Lys and Tyr residues were calculated based on previously established procedures by Zheng et al. and Vernon et al.^7,33^ The Arg/Lys and Tyr interaction was filtered for the distance and angle cutoff. The distance cutoff guarantees the magnitude of the vector joining the charged nitrogen of Arg/Lys and the centre-of-mass of the *π* group in Tyr to be less than 6 Å. Furthermore, the angle cutoff ensures that the absolute cosine of the angle between the previously mentioned vector and the normal vector of the *π* group in Tyr should be greater than 0.8.

The quantifications of sp^2^-*π* interactions between Arg and Tyr are calculated based on the established methodology reported in the recent literature.^7,33^ Firstly, the distance between the centre-of-mass of the Arg guanidinium group and the Tyr benzene ring is constrained within a cutoff of 8 Å. Secondly, the cosine of the angle between the normal vector of the plane defined by the Arg guanidinium group and the Tyr benzene group should be more than 0.8. Finally, both planes defined by the Arg guanidinium and Tyr benzene groups are elevated by 1.5 Å, and the distance between the centre-of-mass of the two new planes is computed. Pairs exhibiting center of mass distances less than 4 Å are identified as forming sp^2^-*π* interactions.

### MUT-16 constructs purification

Three different MUT-16 constructs were purified to test their condensate-formation properties in vitro: MUT-16 633-772 (M8BR), MUT-16 773-944 (FFR), MUT-16 633-944 (M8BR+FFR). The gene fragments coding for MUT-16 (O62011) were cloned into modified pET vectors using ligation-independent cloning. MUT-16 constructs were produced as fusion proteins with an N-terminal MBP-tag and a C-terminal 10xHis-tag in the *E. coli* BL21(DE3) derivatives strain in Terrific Broth (TB) medium. Protein production was induced at 18°C by adding 0.2 mM IPTG for 12-16 hours. Cell pellets expressing the MUT-16 constructs were resuspended in Lysis buffer (20 mM Tris/HCl, 50 mM NaPO4, 500 mM NaCl, 10% (v/v) glycerol, 5 mM 2-mercaptoethanol, pH 7.5) and lysed by sonication. Insoluble material was removed by centrifugation. MUT-16 constructs were purified by immobilised metal affinity chromatography using Ni^2+^-chelating beads (HisTrap FF; GE Healthcare) and eluted with 20 mM Tris/HCl pH 7.5, 150 mM NaCl, 500 mM imidazole, 10% (v/v) glycerol, 5 mM 2-mercaptoethanol. MUT-16 was dialysed against 20 mM Tris/HCl pH 7.5, 150 mM NaCl, 10% (v/v) glycerol, and 5 mM 2-mercaptoethanol. MUT-16 was subsequently purified using anion exchange chromatography (HiTrap Q, GE Healthcare) and size-exclusion chromatography using an S200 increase 16/600 column (GE Healthcare) in a buffer containing 20 mM Tris/HCl pH 7.5, 150 mM NaCl, 10% (v/v) glycerol, 2 mM DTT. All steps were performed on ice or at 4°C. Proteins were stored at −70°C.

### *In vitro* phase separation assays

Purified MUT-16 constructs, MUT-16 633-772 aa (M8BR), MUT-16 773-944 aa (FFR), and MUT-16 633-944 aa (M8BR+FFR), were diluted to a final concentration of 50 µM in the storage buffer, 20 mM Tris/HCl (pH 7.5), 150 mM NaCl, 10% (v/v) glycerol, and 2 mM DTT. A 1:1 serial dilution was performed in 8-strip PCR tubes (Multiply-*µ*strip Pro 8-strip, Sarstedt, REF 72.991.002) for droplet formation. For each construct, the reaction was done in two parallel serial dilutions. The first one, without adding 3C protease, was used as a control. The second serial dilution included the addition of 3C protease (1 mg/mL; in-house produced) at a 1:100 (w/w) ratio of 3C protease to the MUT-16 fragment. 3C protease removed the N-terminal MBP tag from the constructs and induced droplet formation. After incubating the reaction for 60 minutes at room temperature to allow cleaving, the reaction mixture was added to a slide that had been previously attached with 1 cm x 1 cm frames (Thermo Scientific, AB-0576) and then covered with a cover slip 20x20 mm (Roth Karlsruhe). Finally, the slides were imaged using Thunder (Leica), an inverted widefield microscope in bright field mode, with a 100x/1.44 oil lens and a 310 ms exposure time. Images were analysed, and the contrast was adjusted using Fiji/ImageJ (2.14.0/1.54f).

### Co-expression pulldown assays

The gene fragments coding for MUT-8 (Q19672) 1-235 and MUT-16 (O62011) 584-724 were cloned into modified pET vectors using ligation-independent cloning. The MUT-16 (584-724) gene fragments carrying mutations of seven Arginine residues (Arg 642, Arg 657, Arg 658, Arg 682, Arg 689, Arg 698, Arg 699) into either Alanines or Lysines were ordered from Integrated DNA Technologies (IDT). MUT-16 (584-724) constructs were produced as fusion proteins with an N-terminal GST-tag, while MUT-8 (1-235) was produced as fusion proteins with an N-terminal MBP-tag. Vectors carry different antibiotic resistance markers to allow co-expression of MUT-8 and MUT-16 in E. coli. Plasmids containing the genes coding for MUT-16 (584-724) and MUT-8 (1-235) were co-transformed into BL21(DE3) derivative strains to allow co-expression. Cells were grown in 50 mL TB medium shaking at 37°C, and protein production was induced at 18°C by adding 0.2 mM IPTG for 12-16 hours. Cell pellets were resuspended in 4 mL of Lysis buffer (50 mM NaH2PO4, 20 mM Tris/HCl, 250 mM NaCl, 10 mM Imidazole, 10% (v/v) glycerol, 0.05% (v/v) IGEPAL, 5 mM 2-mercaptoethanol pH 7.5). Cells were lysed by sonication, and insoluble material was removed by centrifugation at 21,000xg for 10 minutes at 4°C. 500 µL of the supernatant was applied to 35 µL amylose resin (New England Biolabs) or glutathione resin (Cytiva) and incubated for 1-2 hours at 4°C. Subsequently, the resin was washed thrice with 500 µL of Lysis buffer. The proteins were eluted in 50 µL of Lysis buffer supplemented with 10 mM maltose (amylose resin) or 20 mM of reduced glutathione (glutathione resin), respectively. Input and eluate fractions were analysed using SDS–PAGE and Coomassie staining.

## Results

### Residue-level coarse-grained simulations predict the tendency of MUT-16 IDRs to phase separate *in vitro*

Simulations using the coarse-grained CALVADOS2 model^35,36^ captured the phase-separation propensities of MUT-16’s intrinsically disordered regions (IDRs) in agreement with *in vitro* experiments. They highlighted the molecular determinants of MUT-16 phase behaviour. Previous studies have shown that at ambient temperature, MUT-16 protein cluster *in vivo* and nucleates the formation of *Mutator foci*.^25,29^ Specifically, the C-terminal region of MUT-16 (773-1050 aa) is crucial for MUT-16’s ability to form a scaffold,^29^ while an intermediate region (633-772 aa), termed M8BR, is required for recruiting MUT-8 but is dispensable for MUT-16 foci formation. In this study, we employed the MUT-16 isoform (Uniprot ID: O62011) and focused on understanding the role of its C-terminal in phase separation. We excluded the region encompassing 945-1054 aa from the C-terminal domain due to its tendency to adopt a 3-D structure (Fig. S1A). We first simulated the M8BR+FFR construct (633-944 aa) (Fig. 2A), followed by individual simulations of FFR (773-945 aa) and M8BR (633-772 aa). Both M8BR+FFR and FFR chains exhibited phase separation (Fig. 2A,B), with dilute phase concentration measured at around 0.28 ± 0.04 mM and 0.15 ± 0.06 mM, respectively at 275 K. In contrast, Mut-16 M8BR chains alone did not phase separate spontaneously (Fig. 2C). In agreement with the simulations, the *in vitro* experiments demonstrated the lack of MUT-16 M8BR phase separation at all the concentrations up to 50 µM (Fig. 2C). MUT-16 FFR was observed to phase separate *in vitro* at concentrations 12.5 µM and above, with droplet number and size increasing with concentration (Fig. 2B). Similarly, the MUT-16 M8BR+FFR phase separated at concentrations as low as 6.25 µM, with droplet size increasing, but the number of droplets decreasing, with higher concentrations (Fig. 2A). Furthermore, the timelapse obtained from *in vitro* experiments showed MUT-16 FFR droplets fusing and wetting the coverslip (Fig. S2, Movie S1), which is typical for liquids.^1^ Both simulation and experimental results demonstrate that the FFR segment of MUT-16 is sufficient to drive liquid-liquid phase separation.

**Figure 2:**
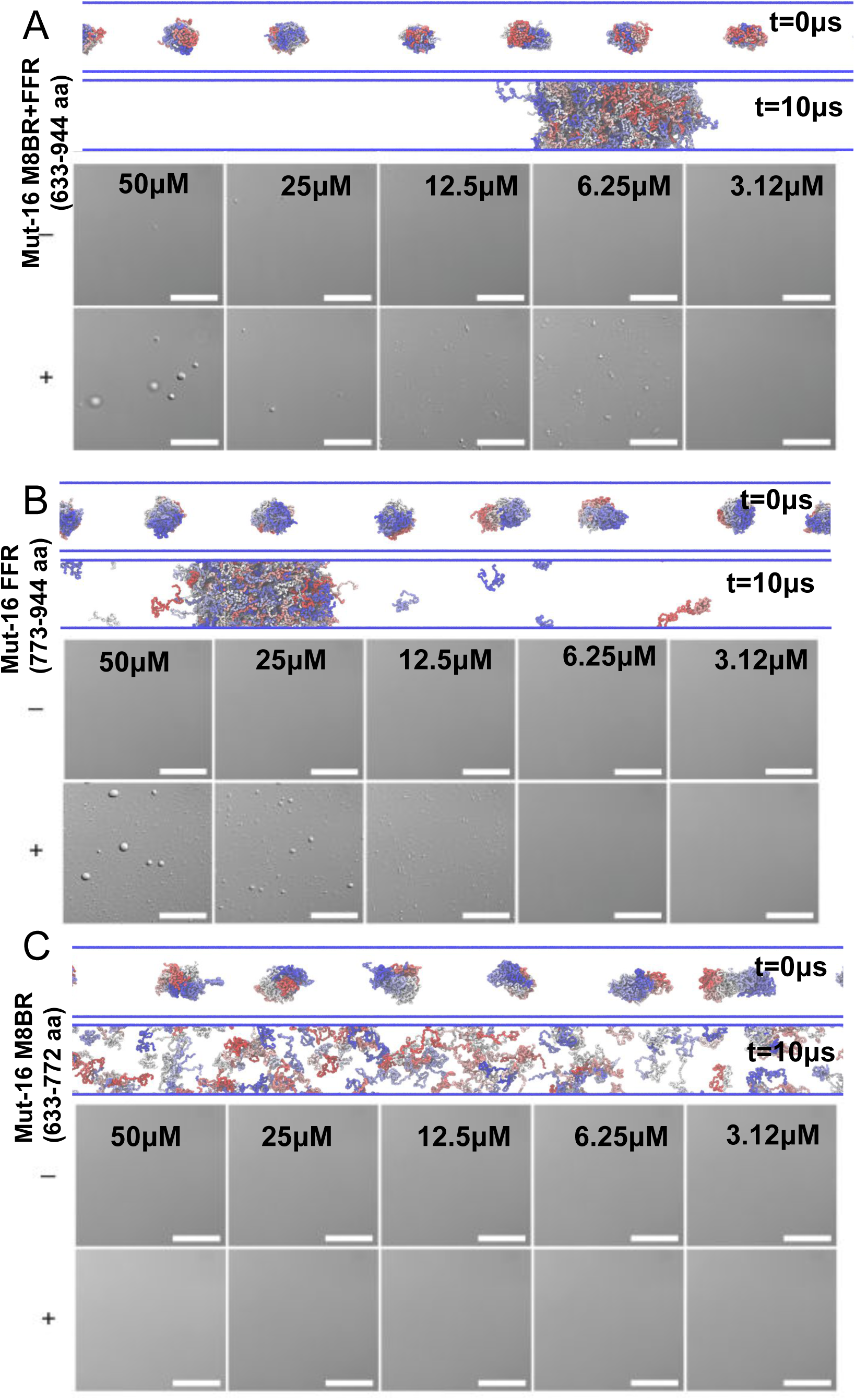
MUT-16 FFR is sufficient to drive the phase separation. **A.** Panel above shows the initial (t=0 µs) and final (t=10 µs) frame of MUT-16 M8BR+FFR simulation in a slab geometry. In the panel below, representative bright-field images show the phase separation assays of the MUT-16 M8BR (633–772 aa) construct as a function of protein concentration. The concentration is indicated at the top of each image. Phase separation was induced by cleaving the MBP N-terminal tag by adding 1:100 (w/w) 3C protease to the MUT-16 fragment. The presence of 3C is represented as +, and its absence is indicated by – on the left part of the panel **B.** Simulation and *in vitro* observations of MUT-16 FFR construct obtained through the method described in **A**. **C.** Simulation and *in vitro* observations of MUT-16 M8BR construct obtained through the method described in **A**. Scale bar correspond to 20 µm.

Further, we performed simulations of MUT-16 M8BR+FFR and FFR at several temperatures between 260 K to 300 K to study the temperature dependence of phase separation. Based on experimental findings from Uebel et al.,,^29^ we expect that MUT-16 M8BR+FFR will cluster together, possibly via phase separation at lower temperatures but not high temperatures. Indeed, at lower temperatures, the MUT-16 M8BR+FFR protein chains phase separates and forms a dense protein-rich phase surrounded by a dilute phase depleted in proteins (Fig. 3A). As the temperature increases, MUT-16 M8BR+FFR chains are progressively observed in the dilute phase (Fig. 3A). This is in qualitative agreement with experiments by Uebel et al.,^29^ where MUT-16 foci disappeared as the worms were exposed to elevated temperature (∼303 K). The condensates reappeared after lowering the ambient temperature (∼294 K). We further established the phase diagram by calculating the concentration of MUT-16 chains in the dense and the dilute phase (Fig. 3B). The phase diagram revealed the critical temperature (T_c_) to be approximately ∼296 K. The agreement of simulations with a transferable physics-based model^30,34,35^ with *in vivo* behaviour adds theoretical support to the interpretation of the temperature-dependent loss of MUT-16 foci in *C. elegans* as upper critical solution temperature (UCST) phases separation.^6^ We also constructed a phase diagram for FFR from CALVADOS2 simulations, with T_c_) of approximately ∼300 for FFR, which is similar to T_c_ for MUT-16 M8BR+FFR, considering that accurate determination of T_c_ can be challenging. While the phase diagram further shows that concentration in the dilute phase is lower for FFR than M8BR+FFR, the comparison demonstrates that the FFR region on its own can phase separate nearly as well as M8BR+FFR (Fig. 3B).

**Figure 3:**
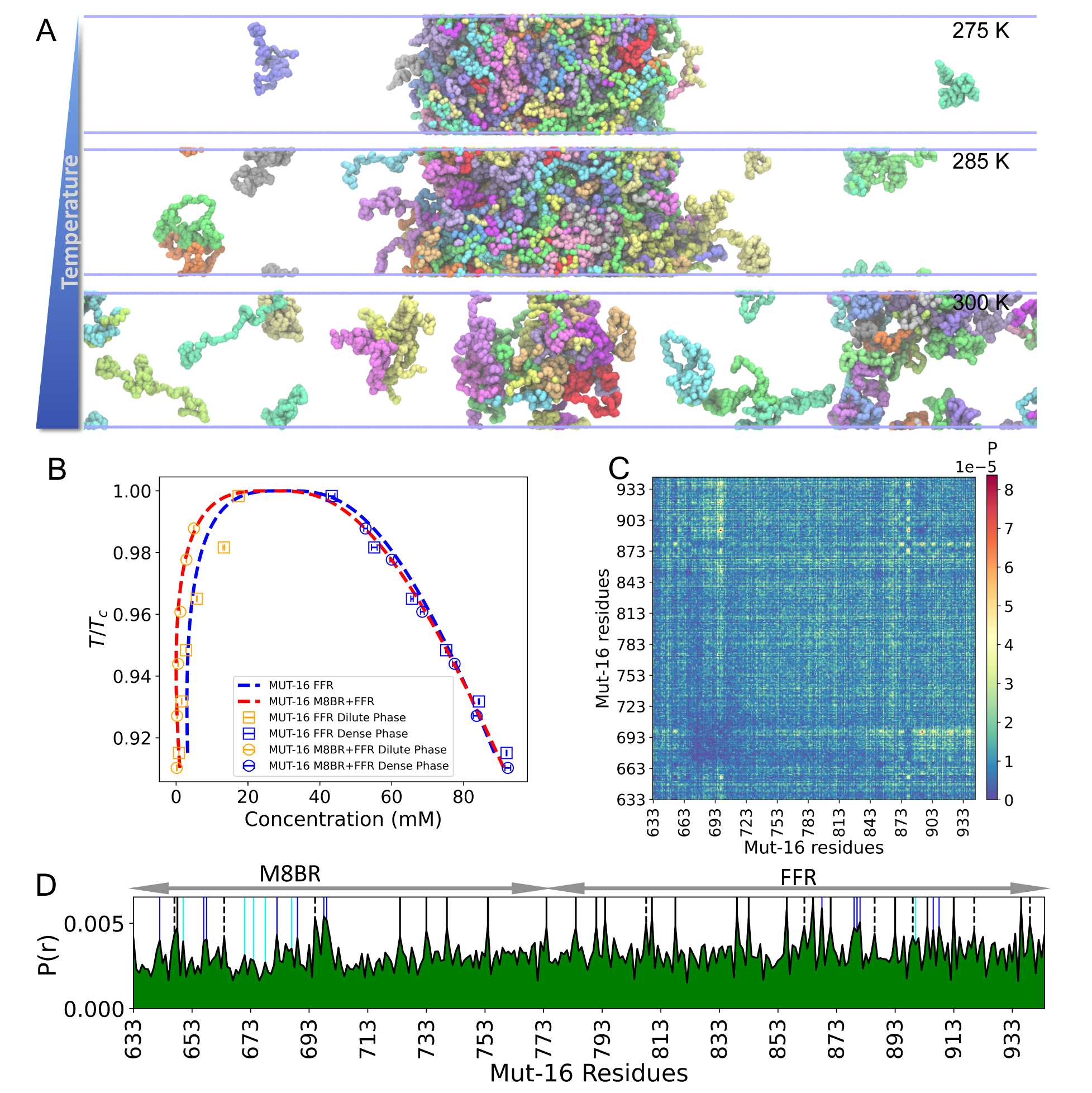
Phase behaviour of MUT-16 condensate in residue-level coarse-grained (CALVA-DOS2) simulations in implicit solvent. **A.** Simulations of 100 MUT-16 M8BR+FFR chains at different temperatures (275 K, 285 K, and 300 K). **B.** Phase diagram of MUT-16 FFR and Mut-16 M8BR+FFR obtained by CALVADOS2 simulation at different temperatures. Blue and red dots represent the dilute and dense phases concentrations of MUT-16 FFR and Mut-16 M8BR+FFR, respectively. **C.** 2-D inter-chain contact-map obtained from the simulation of MUT-16 M8BR+FFR chains at *T /T_c_* = 0.92. **D.** One-dimensional contact map representing the peaks associated with the amino acid with higher relative contact probabilities (P(r)). The peaks shown by Tyr (black), Phe (dashed), Arg (blue), and Lys (cyan) are represented by the vertical lines.

CALVADOS2 simulations further identified key regions and amino acids driving the phase separation of MUT-16. Inter-molecular contact probabilities obtained from the CALVADOS2 simulation of MUT-16 M8BR+FFR at 0.92 T/T_c_ revealed that the FFR region exhibits a stronger propensity for interaction compared to the M8BR (Fig. 3C). While residues throughout the sequences contribute to interactions (Fig. 3C), the region spanning residues 663 to 693 and residues 703 to 713 in M8BR exhibits notably fewer interactions. A region-wise contact map demonstrated a higher FFR:FFR interaction probability followed by FFR:M8BR and M8BR:M8BR (Fig. S3A,B), suggesting a correlation with the high phase separation propensity of FFR observed in both CALVADOS2 simulations and experiments. Although the M8BR has a lower interaction probability, it maintains substantial contact with the FFR. The 1-D contact map highlighted the increased interaction probability of aromatic residues(Fig. 3D), with Tyr showing the most prominent and abundant peaks, followed by Phe. The abundance of aromatic residues in FFR (14%) compared to the M8BR (5%), likely enables the FFR to drive the phase separation. Additionally, Arg residues showed elevated interaction propensities, with Arg 698 and Arg 699 forming substantial contacts (Fig. 3D). In contrast, Lys residues displayed a lower tendency to interact, with a stretch of Lys in M8BR (Lys 671, Lsy 674, and Lys 678) showing only small spikes in contact probability (Fig. 3D). The presence of significant Tyr interaction peaks in the M8BR region (700-772 aa) aligns with experimental findings by Uebel et al.,^29^ where a GFP-tagged MUT-16 construct (704-1050 aa) forms larger foci compared to a shorter MUT-16 construct (773-1050 aa). Further, we computed the interaction frequency of amino acid pairs normalised (Fig. S4A) and unnormalised (Fig. S4B). The normalized contact map reveals residue-residue interactions, ranking the pairs with the highest interaction probability as follows: Tyr-Arg, Tyr-Tyr, Tyr-Phe, Phe-Arg, Phe-Phe, Arg-Asp, Arg-Glu, and Cys-Cys (Fig. S4A). Additionally, somewhat weaker interactions were observed between Tyr and Phe with residues like Leu, Ile, His, Met, Cys, Gln, and Asn. The unnormalised interactions also reveal the drivers of phase separation, as many weak interactions can be equally or more important than a more limited number of relatively strong interactions.^11^ The unnormalized contact map indicates that Pro-Pro, Pro-Gln, Gln-Gln, Pro-Tyr,^65^ Gln-Tyr, Pro-Asn, and Gln-Asn interactions are the most abundant, listed in decreasing order of interaction probability(Fig. S4B). While some residue pairs dominate, a broad range of residues also contributes to phase separation.^11^ Overall, our simulations demonstrate the critical role of MUT-16 FFR in phase separation and reveal the specific amino acids involved in this process.

### MUT-16 phase separation in explicit solvent coarse-grained Martini3 simulations

To better characterise the molecular drivers of MUT-16 phase behaviour, we performed near-atomic resolution explicit-solvent Martini3 coarse-grained simulations.^39,40^ Martini3 is a powerful general-purpose coarse-grained model of biomolecules that have been successfully used to simulate multi-domain and disordered proteins.^40–43^ However, unlike CALVADOS2, Martini3 has not been extensively tested for phase separation in disordered proteins.^40^ Therefore, comparing Martini3 simulations with other models, such as CALVADOS2 or atomistic simulations, as well as experimental data, is essential for validation and consistency. In our Martini3 simulations, we observed the spontaneous phase separation of the MUT-16 M8BR+FFR (Movie S2), MUT-16 FFR and MUT-16 M8BR chains suspended in explicit solvent. Since the Martini3 force-field is known to overestimate protein-protein interactions, we re-scaled the protein-water interactions using a rescaling factor *λ*.^42,66^ We reasoned that if the propensity of MUT-16 M8BR+FFR, MUT-16 FFR and MUT-16 M8BR phase separation, captured by Martini3, results from favourable interactions in the simulations, then increasing protein-water interactions somewhat should not abrogate condensation. At lower *λ* values of 1.00 and 1.01, condensates of all three constructs (MUT-16 M8BR+FFR, MUT-16 FFR and MUT-16 M8BR) persisted (Fig. 4A,B,C), as indicated by the density profiles. At larger *λ* values (1.05), all three condensates dissolved(Fig. 4A,B,C). However, at an intermediate lambda value of 1.03, the MUT-16 M8BR+FFR and MUT-16 FFR condensates persist, but MUT-16 M8BR condensate dissolves (Fig. 4D,E,F).

**Figure 4:**
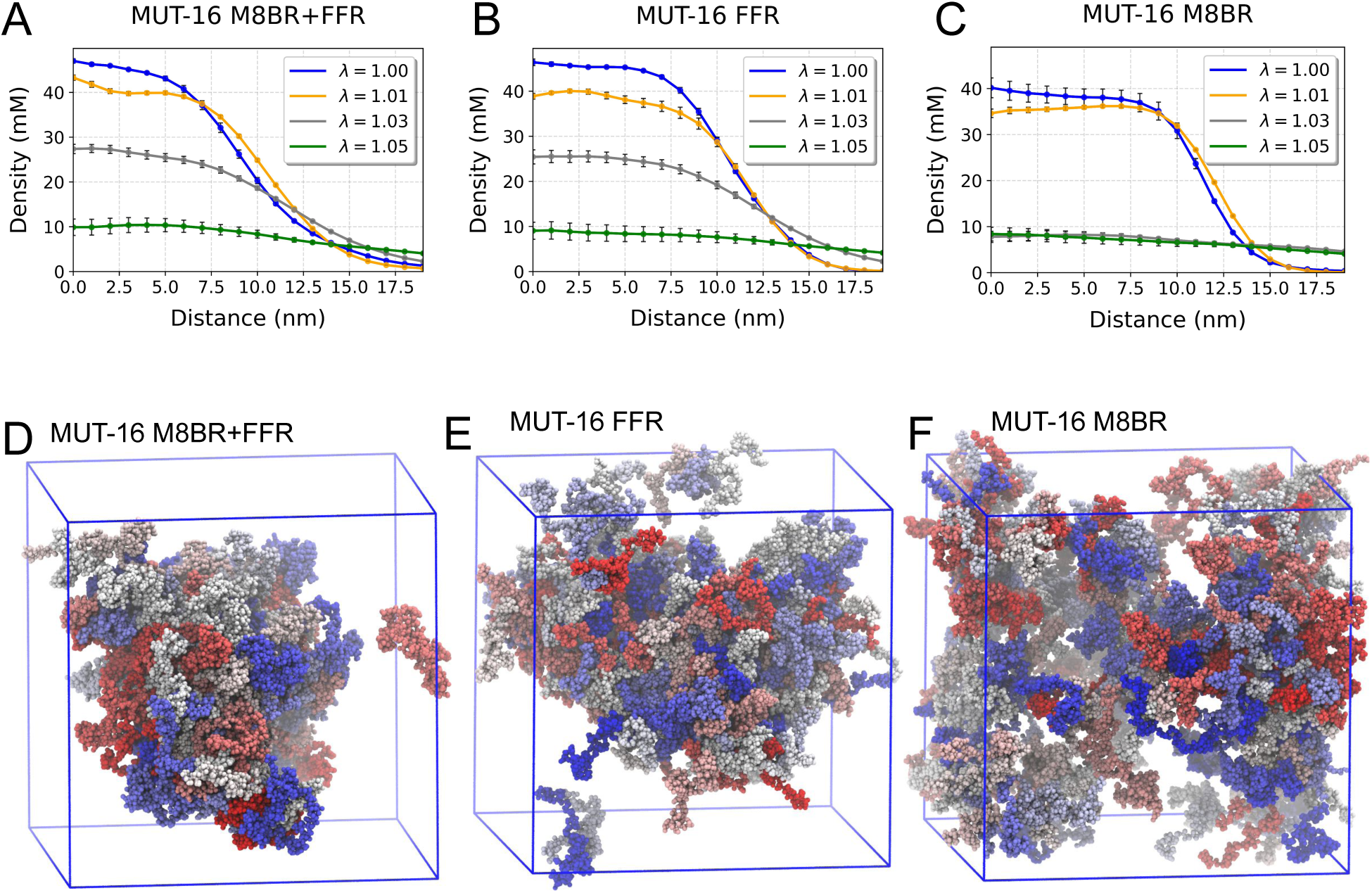
Martini3 simulations of MUT-16 constructs at various *λ* values (*λ* = 1, 1.01, 1.03, 1.05). **A,B,C.** Density profiles of MUT-16 M8BR+FFR, MUT-16 FFR, and MUT-16 M8BR. The x-axis denotes the distance (nm) from the protein’s centre-of-mass, while the y-axis indicates the protein density (mM). At *λ* = 1.03 (grey), there is a significant difference in the density profiles among the constructs, indicating phase separation of MUT-16 M8BR+FFR and MUT-16 FFR, but not MUT-16 M8BR. **D,E,F.** Final snapshot of 10 µs Martini3 simulations of MUT-16 M8BR+FFR, MUT-16 FFR and MUT-16 M8BR constructs at *λ*=1.03.

We further characterized the regions and specific residues within MUT-16 M8BR+FFR that drive phase separation using a contact map derived from the Martini3 simulation at *λ* = 1.03 (Fig. 5A). The 2-D contact map revealed a concentration of contacts within MUT-16 FFR(Fig. 5A), consistent with our CALVADOS2 simulations and *in vitro* experiments. Interestingly, a sub-region of MUT-16 (850-940aa) was identified as particularly critical for phase separation. The 1-D contact map highlighted the peaks corresponding to aromatic (Tyr, Phe) and positively charged residues (Arg, Lys) (Fig. 5C), suggesting their involvement in phase separation. Interestingly, the region-wise contact map from Martini3 showed that FFR:FFR interaction probability was not significantly more substantial than M8BR:M8BR (Fig. S3B), a result contrasting with the CALVADOS2 simulations (Fig. S3A), where FFR:FFR interactions were more prominent. Further, the 2-D contact map revealed stronger interaction between the N-terminal region (640-700aa) and the C-terminal region (701-773aa) of MUT-16 M8BR (Fig. 5A), which were absent in the CALVADOS2 simulations (Fig. 3C). A comparison of the 1-D contact map from Martini3 and CALVADOS2 simulations highlights a higher contribution of MUT-16 M8BR N-terminal region (640-700aa) in Martini3 simulations compared to the CALVADOS2 (Fig. S3C). Unlike CALVADAOS2, Martini3 showed elevated contact probabilities of Lys residues (Lys 671, Lys 674, and Lys 678)(Fig. S3C), suggesting that Lys residues are probably more “sticky” in Martini3 than in CALVADOS2.

**Figure 5:**
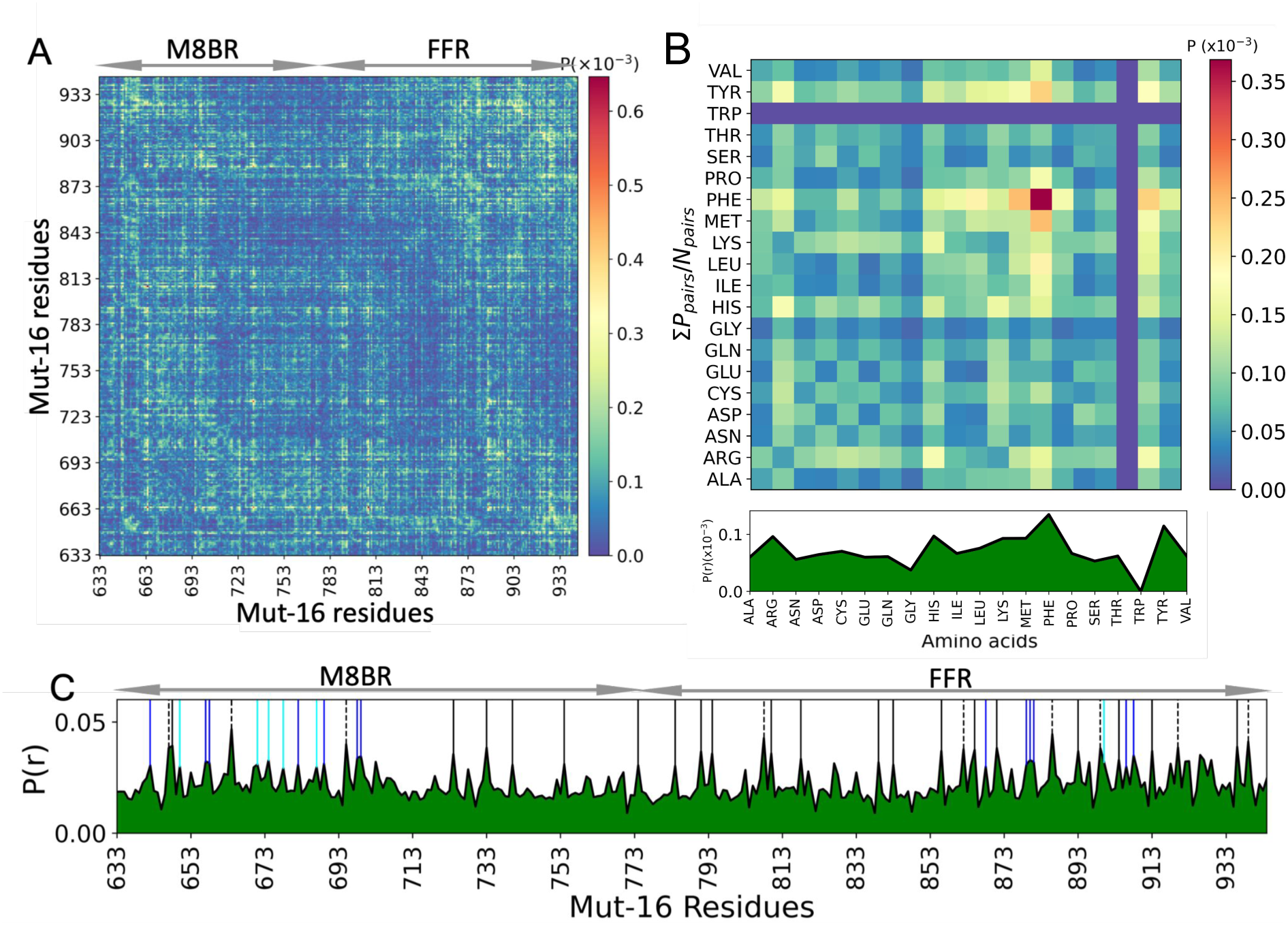
MUT-16 phase separation in explicit-solvent coarse-grained (Martini3) simulations. **A.** Inter-molecular contact probabilities in the condensed phase of MUT-16 M8BR+FFR as a function of the amino acid sequence. The MUT-16 M8BR and FFR regions are highlighted. **B.** Inter-molecular contact probability of MUT-16 M8BR+FFR as a function of residue type normalised by the relative abundance of amino acid pairs. **C.** One-dimensional contact map representing the peaks associated with the amino acid with higher relative contact probabilities (P(r)). The peaks shown by Tyr (black), Phe (dashed), Arg (blue), and Lys (cyan) are represented by the vertical lines.

Additionally, we computed normalised and unnormalised amino acid-based contact maps to examine the contributions of specific residues to phase separation. In the normalised contact map, aromatic and positively charged residues, such as Tyr, Phe, Arg, Lys and His, played crucial roles in the phase separation of MUT-16 (Fig. 5B). Interestingly, Phe-Phe interactions were stronger than Tyr-Tyr interactions. Furthermore, the unnormalised amino-acid-based contact map (Fig. S5) indicates a higher propensity of Pro-Pro and Pro-Tyr^65^ interactions, followed by somewhat weaker interactions involving His (e.g., His-Pro), Gln (e.g., Gln-Gln), and Asn (e.g., Asn-Gln) as shown by the sum of the number of contacts in Fig. **??**. Tyr has a higher overall contribution in phase separation than Phe due to its abundance and interaction strength. In conclusion, the interaction propensities observed in Martini3 simulations were broadly comparable to those seen in CALVADOS2 simulations^36^ (Fig. S3C), with the key exception of Lys residues. As CALVADOS2 was specifically parameterised to capture the interactions of amino acids in IDRs and IDP conformations,^35,36^ the overestimation of Lys interactions in Martini3 simulations likely reflects a limitation in the force field’s ability to model IDR and IDP phase separation accurately.

### Recruitment of MUT-8 N-terminal PLD by MUT-16 condensate

With simulations at two different levels of resolutions (residue-level CALVADOS2 and near-atomic resolution Martini3) providing a consistent view of MUT-16 phase separation, we extended our investigation to examine the role of MUT-16 as a scaffold for recruiting its client protein, MUT-8 (Uniprot ID: Q58AA4). Interestingly, sequence analysis revealed that the first 51aa of MUT-8 is predicted to be a prion-like domain (PLD), Fig. S1),^67^ leading us to hypothesize that this region could interact with MUT-16 condensates. In simulations with both models, the MUT-16 M8BR+FFR condensate remained stable and spontaneously recruited the MUT-8 N-terminal region (Fig. 6A,B). Further, analysis of contact maps from CALVADOS2 (Fig. S6C) and Martini3 (Fig. 6C) simulations revealed key regions and residues involved in the interaction between MUT-16 M8BR+FFR and MUT-8 N-terminal domain. Two significant findings emerged: (i) In CALVADOS2 simulations, MUT-8 recruitment is primarily facilitated by the MUT-16 FFR (Fig. S6A) (ii) In the Martini3 simulations, recruitment is predominantly driven by the MUT-16 M8BR (Fig. S6B), with a smaller contribution from MUT-16 FFR. Both models suggest that recruitment is facilitated by both the M8BR and FFR regions (Fig. 6C,S6D). The *in vivo* experiments show loss of MUT-8 recruitment upon deletion of the M8BR (633-772 aa) region and loss of foci formation upon deletion of a region (773-1050 aa) including the FFR. The experiments suggest that the M8BR is essential for MUT-8 recruitment but do not rule out that the FFR region further enhances MUT-8 recruitment. Interestingly, we observed that the positively charged residues (Arg and Lys) in the MUT-16 M8BR (Fig. 6C, Fig. S6D) interact with the Tyr residues of MUT-8 N-terminal domain (Fig. S6D), possibly through cation-*π* or sp^2^-*π* interactions. Similarly, the aromatic residues (Tyr and Phe) in MUT-16 FFR interact with Tyr residues of MUT-8 N-terminal domain(Fig. 6C,Fig. S6D), likely via *π*-*π* interactions. A comparison of the 1-D contact maps revealed a lower contact probability of Lys residues in MUT-16 M8BR in the CALVADOS2 model compared to the Martini3 model (Fig. S6D), further supporting the hypothesis that Lys residues exhibit increased “stickiness” in Martini3.

**Figure 6:**
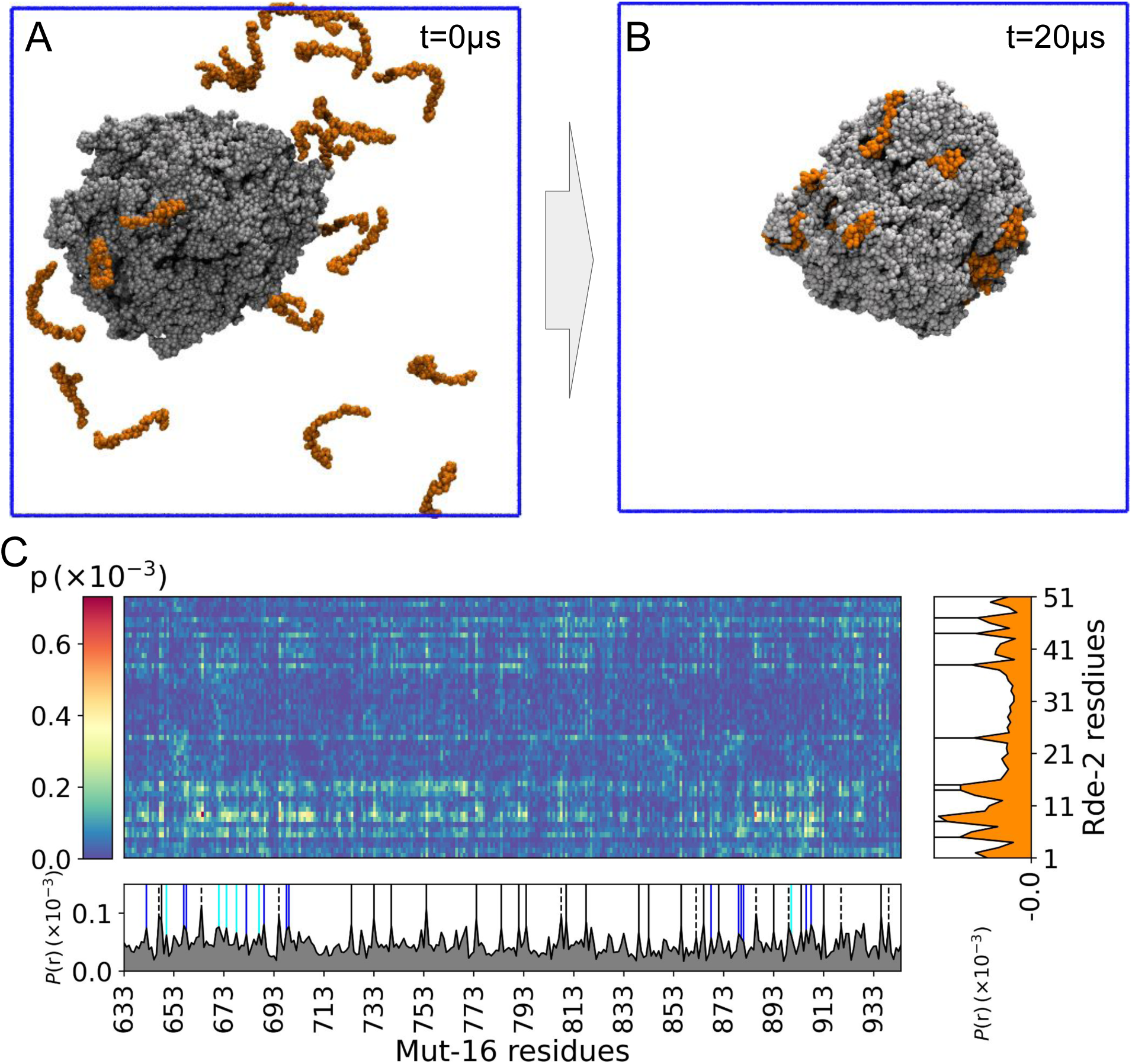
Recruitment of MUT-8 N-terminal PLD driven by phase-separated MUT-16 M8BR+FFR. **A,B.** Initial and final snapshots of a 20 µs Martini3 simulation of the interaction between MUT-8 N-terminal PLD (orange) and MUT-16 M8BR+FFR (grey) phase-separated condensate. **C.** Inter-molecular contact probabilities of interaction between amino acids of MUT-8 N-terminal and MUT-16 M8BR+FFR are shown in a 2-D contact map. 1-D relative contact probability (P(r)) of the MUT-16 (grey) and MUT-8 (orange) amino acids is plotted to indicate the higher peaks by positively charged (Arg (blue), Lys (cyan)) and aromatic (Tyr (black), Phe (dashed)).

Given that Uebel et al.^29^ demonstrated a loss of MUT-8 recruitment *in vivo* upon deletion of the MUT-16 M8BR, we sought to investigate the recruitment mechanism of MUT-8 N-terminal by MUT-16 M8BR. We designed a minimal simulation setup to disentangle MUT-8 recruitment from the phase separation of MUT-16 *in silico*. In this setup, we performed Martini3 simulations MUT-16 M8BR and MUT-8 N-terminal domain at *λ* = 1, simulating conditions where altered solution conditions, such as the presence of a crowding agent stabilise an otherwise unstable M8BR condensate. In this simulation, we observed the clustering of the MUT-16 M8BR chains and subsequent recruitment of the MUT-8 N-terminal domain (Fig. S7A). Upon increasing the *λ* to 1.03, the MUT-16 M8BR cluster bound to MUT-8 starts to dissolve(Fig. S7B). Interestingly, the contact map indicated an essential role of the very N-terminal residues of MUT-16 M8BR (640-700aa) in the recruitment mechanism (Fig. S7C). Furthermore, it highlighted the importance of interaction between positively charged residues (Arg and Lys) of MUT-16 M8BR and Tyr of MUT-8 N-terminal domain (Fig. S7C), further supporting the potential involvement of cation-*π* or sp^2^-*π* interaction in MUT-8 recruitment. We quantified the cation-*π* interactions between ARG/LYS of MUT-16 M8BR and TYR of MUT-8 N-terminal domain^33^ and observed that Arg exhibits more frequent cation-*π* interactions with Tyr compared to Lys residues (Fig. S8) in line with previous experiments.^10^ However, it is essential to highlight that Martini3 simulations may lack the chemical detail necessary to fully capture the complexity of non-covalent interactions such as cation-*π*, sp^2^-*π* and hydrogen bonding.

### Atomistic molecular dynamics simulations reveal the role of cation-*π*, sp^2^-*π*, and hydrogen bond in recruitment of MUT-8 N-terminal PLD

We performed atomistic molecular dynamics simulations in explicit solvent to investigate the specific interactions between MUT-16 M8BR and the MUT8 N-terminal domain responsible for its recognition. While the overall timescales involved in molecular recognition exceed the current capabilities of atomistic simulations, we aimed to elucidate residue-residue contacts, which frequently break and reform within the microsecond timescale accessible to atomistic simulations.^32,33,68^ The atomistic simulations (Fig. 7A) were initiated by backmapping the final frame of a 20 µs Martini3 coarse-grained slab simulation.^58^ This backmapping converted the coarse-grained system to an atomistic representation containing ≈ 900, 000 atoms, which was then simulated for 1 µs. Further, we extracted 100 equidistant frames from the 1 µs simulation and launched 100 independent simulations from each frame, totalling up to 350 µs of sampling on Folding@home.^64^

**Figure 7:**
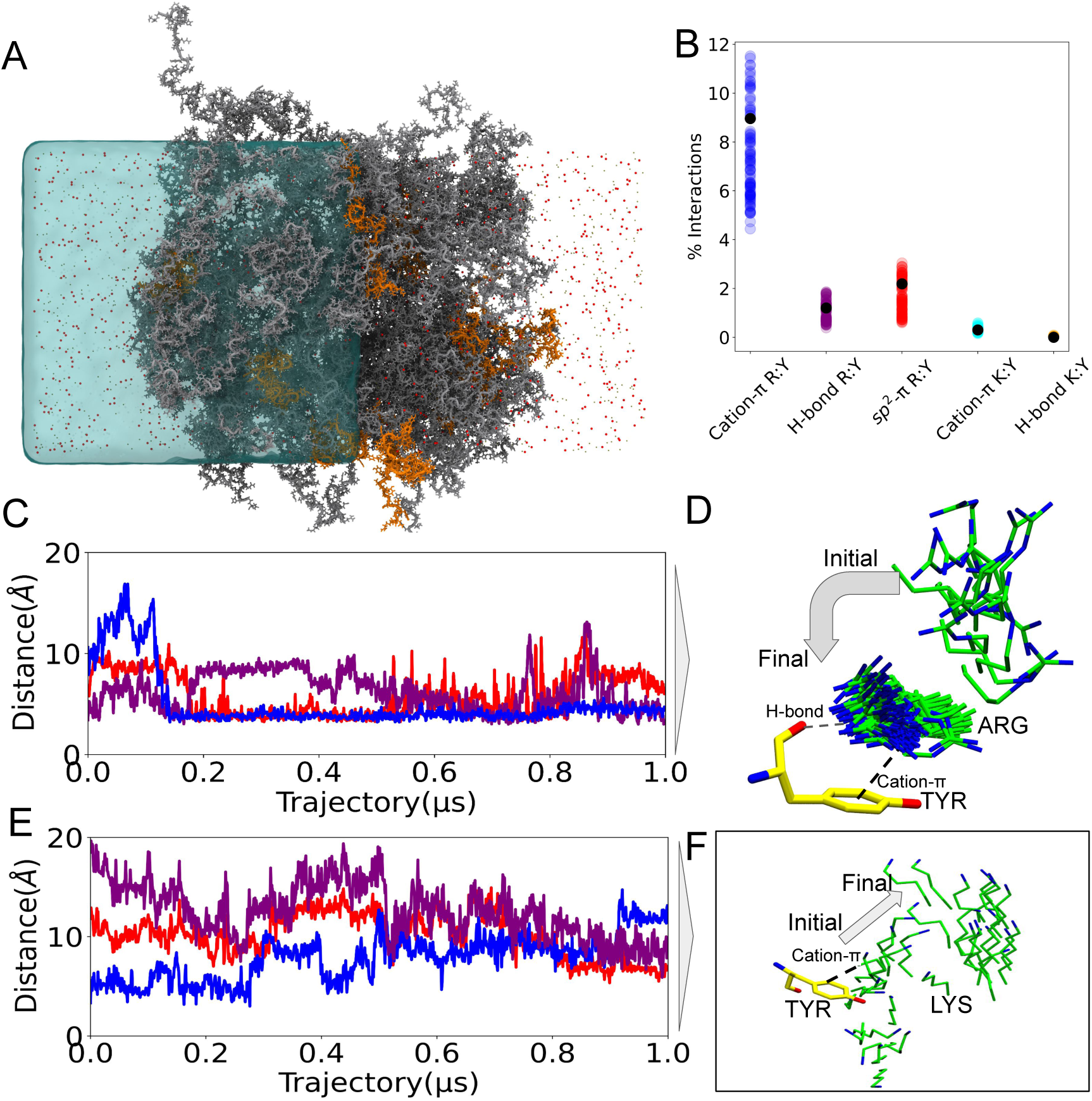
**A.** Atomistic simulation depicting 100 chains of MUT-16 M8BR (grey) and the N-terminal domain of MUT-8 (orange). Sodium (*Na*^+^) and chloride (*cl^−^*) ions are shown as olive and red points, respectively, suspended in water (blue). For clarity, the water on the right half of the simulation box is deleted. **B.** Comparison of the percentage of cation-*π*, sp^2^-*π* and H-bond interactions made by Arg/Lys (MUT-16 M8BR) and Tyr (MUT-8 N-terminal). The percentage of H-bonds was calculated given the pair was already forming the cation-*π* interaction. Multiple data points were calculated from the 100 atomistic runs obtained from Folding@home (coloured, translucent) and in-house 1 µs trajectory (black, opaque). **C.** Time evolution of the distance between the aromatic ring of Tyr and guanidinium group of Arg for selected pairs of residues. **D.** The pictorial representation of one of the residue pairs whose time evolution (blue) is plotted in **C**. **E.** Time evolution of the distance between the aromatic ring of Tyr and the amino group of Lys for selected amino acid pairs. **F.** The pictorial representation of one of the residue pairs whose time evolution (blue) is plotted in **E**.

The contact map obtained from the 350 µs atomistic simulation was roughly analogous to the results from Martini3 simulations (Fig. S9A), validating the role of Arg-Tyr interaction in MUT-8 recruitment. However, Lys residues showed a lower interaction probability compared to Martini3, further indicating that they may be overly prone to interactions in the context of IDR-IDR interactions within the Martini3 force field. To investigate the atomistic system in more detail, we quantified the stability of hydration water bound through hydrogen bonds to the amino-acids, as well as the behaviour of ions interacting with charged residues throughout 1 µs in-house atomistic trajectory (Fig. S9B). Hydration water remained stable over time, but the number of bound ions decreased over time, with Na^+^ fluctuating more than Cl*^−^* ions. Further, we quantified the density of water, MUT-16 M8BR, MUT-8 N-terminal domain, and ions over the 350 µs trajectory. The dense-phase water density, ≈ 600 mg/ml (Fig. S9C), was consistent with previous studies on FUS low-complexity region.^33,69^ The density of MUT-16 M8BR was ≈ 600 mg/ml, while MUT-8 N-terminal density was around ≈ 20 mg/ml in the dense phase (Fig. S9C). Furthermore, we observed higher molarity of Na^+^ and Cl*^−^* ions in the dilute phase than in the MUT-16 M8BR cluster (Fig. S9D). Zheng et al.^33^ proposed a method to rationalise how ions are distributed in simulations of protein condensates based on the local concentrations of cationic and anionic residues in a condensate. With this approach, we predicted concentration profiles of Na^+^ and Cl*^−^* ions and found that overall the predicted concentrations (dashed lines) matched the concentration profiles from the simulations (Fig. S9D). There are more Cl*^−^* ions than Na^+^ ions in MUT-16 M8BR cluster due to the multiple positively charged residues (Arg and Lys) present in the N-terminal region of MUT-16 M8BR.

To further investigate the system we quantified the contribution of cation-*π* or sp^2^-*π* interactions in Arg-Tyr and Lys-Tyr pairings (Fig. 7B). We tracked the dynamics of these interactions between MUT-8 and MUT-16 throughout the 1 µs initial trajectory and subsequent 350 µs simulations on Folding@Home.^64^ The abundance of cation-*π* and sp^2^-*π* interactions were calculated following the protocols established in previous studies^33^ (see Methods). As anticipated, Arg in MUT-16 exhibited a higher frequency of cation-*π* interactions with Tyr in MUT-8 compared to Lys in MUT-16 (Fig. 7B). We also observed that the Arg residues form sp^2^-*π* interactions with Tyr (Fig. 7B) due to the planar sp^2^ hybridised guanidinium group in Arg. In contrast, Lys cannot form these interactions with Tyr because its side chain contains a flexible primary amine group, which lacks the necessary planar structure. We analysed Arg-Tyr and Lys-Tyr interactions by measuring the distance between the guanidinium group of Arg (or the amino group of Lys) and the center-of-mass of Tyr’s benzene ring (Fig. 7C,E). We observed both persistent interactions, stable for hundreds of nanoseconds, and transient interactions that broke within just a few nanoseconds. Persistent Arg-Tyr contacts (blue line) were established within 150 ns and maintained throughout the trajectory (Fig. 7C). The conformation of this Arg group relative to the Tyr in MUT-8, shown in Fig. 7D, reveals that the guanidinium group aligns parallel to the aromatic ring of Tyr. In contrast, other Arg-Tyr pairs exhibited transient contacts (red and purple lines), with contact durations of less than 100 ns Fig. 7C. Lys-Tyr interactions appeared less stable. Fig. 7E shows the Lys-Tyr pair maintaining contact for about 250 ns (blue line) before dissociating. Conformations of the Lys making the initial contact and conformations after this Lys-Tyr contact is broken are shown in Fig. 7F. Other Lys-Tyr pairs (red and purple lines) formed close interactions only toward the end of the simulation (Fig. 7E). Nevertheless, these contacts were weaker and less frequent than Arg-Tyr interactions, highlighting that, for these exemplary contact pairs, Arg forms more interactions than Lys, which are also less transient.

Visualisation of simulations illustrated that Arg can form a hydrogen bond with the backbone carbonyl of Tyr in addition to cation-*π* and sp^2^-*π* interactions (Fig. 7D). Furthermore, we also observed the coexistence of cation-*π*, sp^2^-*π* and hydrogen bond in Arg and Tyr pairs (Fig. S10), with the three interactions predominately coinciding for an exemplary Arg-Tyr pair, except for short transients, e.g., shortly after 0.4 µs when a sp^2^-*π* interaction is detected but no cation-*π* interaction or hydrogen bond. Unlike Arg, Lys does not form additional hydrogen bonds with the Tyr backbone when involved in cation-*π*(Fig. 7B), limiting the stability of these interactions. The presence of this “triple” interaction profile comprising cation-*π*, sp^2^-*π* and hydrogen bond may explain the robustness and stability of Arg-Tyr interactions compared to Lys-Tyr interactions. This finding is in line with previous quantum chemical calculations^15,70,71^ as well as by a recent study involving an artificial IDP,^11^ where atomistic molecular dynamics simulations of condensates also resolved interactions involving hydrogen bonds with the backbone. These observations highlight the critical role of Arg residues in Mut-16 M8BR and Tyr in MUT8 N-terminal PLD in driving the recruitment of MUT-8 to the *Mutator foci*.

### Co-expression pulldown experiments validate the importance of Arg residues in M8BR of MUT-16 for MUT-8 recuitment

To further investigate the role of Arg in Mut-16 M8BR in the recruitment mechanism of MUT-8, we performed co-expression pulldown experiments using a wt MUT-16 fragment (584-724), which contains the N-terminal of MUT-16 M8BR, and MUT-8 N-terminal (1-235), which includes the prion-like domain (Fig.S1). In addition, we performed similar pulldown experiments with MUT-16 variants where seven key Arg residues within M8BR (Arg642, Arg 657, Arg 658, Arg 682, Arg 689, Arg 698, Arg 699) were mutated to either Lys (Arg *>* Lys) or Ala(Arg *>* Ala) residues. The Arg residues for mutation were selected based on their high interaction frequency in the contact maps obtained from the atomistic (Fig. S9A) and Martini3 (Fig. S7C) simulations. The results demonstrated a reduced binding affinity between MUT-16 Lysine mutants (Arg *>* Lys) and MUT-8 N-terminal compared to wt MUT-16 (Fig. 8). Furthermore, the MUT-16 alanine mutant (Arg *>* Ala) exhibited an even more pronounced decrease in binding affinity, indicating a further loss of MUT-8 recruitment (Fig. 8). This trend was consistent regardless of whether an MBP-tagged MUT-8 N-terminal fragment or a GST-tagged MUT-16 fragment was used for the pulldown. These findings strongly reinforce the results from both coarse-grained and atomistic simulations, demonstrating that Arg residues in MUT-16 M8BR play an essential role in recruiting MUT-8 to the *Mutator foci*.

**Figure 8:**
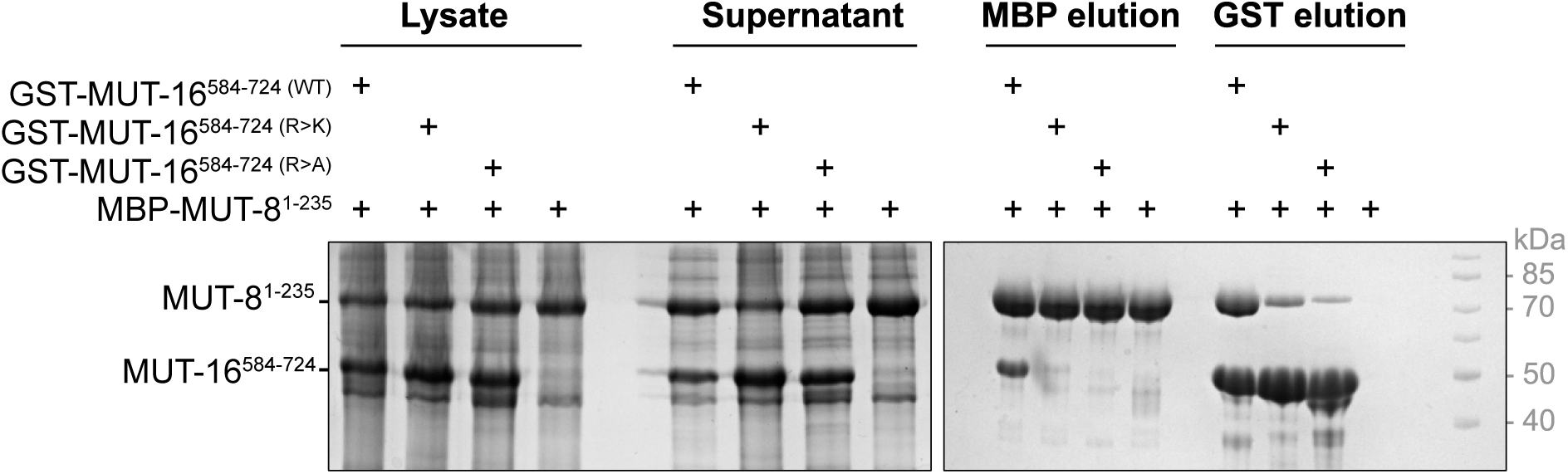
Contribution of Arg residues of MUT-16 M8BR in MUT-8 recruitment. Analysis of the interaction between an MBP-tagged MUT-8 N-terminal fragment (1-235) and a GST-tagged MUT-16 fragment (584-724) by pulldown assays. Three different MUT-16 constructs are tested: wild-type, a construct where seven Arg residues (642, 657, 658, 682, 689, 698, 699) were mutated to Lys (R *>* K), and a construct where the identical seven Arg residues were mutated to Ala (R *>* A). MUT-16 constructs were co-expressed with MUT-8 in *E. coli*. Both MBP and GST pulldowns were performed. SDS-PAGE, followed by Coomassie staining, analysed the total Lysate, supernatant, and elution.

## Discussion

MUT-16 is the scaffolding protein of *Mutator foci* in *C. elegans* germ cells, playing a crucial role in RNA silencing. It recruits several binding partners to *Mutator foci*, including MUT-8, which further recruits the exoribonuclease MUT-7. In this study, we employed a multiscale approach combining coarse-grained simulations (residue-level CALVADOS2 and near-atomic Martini3), atomistic simulations, and *in vitro* experiments to elucidate how MUT-16 undergoes phase separation to form *Mutator foci* and recruits the MUT-8 N-terminal PLD.

Our coarse-grained simulations and *in vitro* experiments show that MUT-16 condensates form through phase separation. CALVADOS2 and Martini3 simulations successfully captured the phase separation tendencies of the disordered regions (M8BR, FFR, and M8BR+FFR). The simulations showed that while MUT-16 M8BR+FFR and MUT-16 FFR spontaneously phase separate, MUT-16 M8BR does not, aligning with our *in vitro* results. The apparent temperature dependence of MUT-16 phase separation in simulations aligns with the temperature-dependent dissolution of MUT-16 foci in *C. elegans*. However, modifications to the residue-level coarse-grained model are required to study whether high or low temperatures favour phase separation. In simulations with the Martini3 model, we found that increased protein-water interactions destabilised the condensate.^42,66^ An increase in protein-water interactions could be interpreted as a change in effective temperature.^44^ Both coarse-grained models emphasise the FFR region as critical for phase separation, highlighting the significant inter-molecular interactions of amino acids such as Tyr, Arg, Phe, Pro, Gln, and Asn in the phase-separated protein condensates. Unlike CALVADOS2, Martini3 reveals stronger interaction involving the M8BR region, comparable to the FFR. This difference is primarily driven by the higher interaction propensity of Lys residues in M8BR, which is probably more “sticky” in Martini3 than CALVADOS2. Additionally, Martini3 simulations show a stronger interaction propensity for Phe residues compared to Tyr in the normalised amino acid-wise contact map. In contrast, CALVADOS2 simulations highlight Tyr as making stronger interactions. Furthermore, the unnormalized amino acid-wise contact map indicates a lower interaction propensity for Gln residues in Martini3 (Fig. **??**) compared to CALVA-DOS2 simulations (Fig.**??**B).

Previous studies have demonstrated that MUT-8 interacts with MUT-16^29^ and that its C-terminus is responsible for recruiting MUT-7.^28^ However, the exact molecular mechanism by which MUT-16 specifically recruits MUT-8 remains elusive. Interestingly, the N-terminus of MUT-8 contains an intrinsically disordered region (IDR), predicted to be a prion-like domain enriched with residues commonly associated with phase separation. To investigate the recruitment of the MUT-8 N-terminal domain to the MUT-16 M8BR+FFR condensate, we performed simulations using both the Martini3 and CALVADOS2 models. Results from both models indicate that interactions with both the M8BR and FFR regions of MUT-16 facilitate the recruitment of the MUT-8 N-terminal domain. Additionally, the contact maps revealed that the MUT-8 N-terminal domain predominantly interacts with Arg-Tyr in MUT-16 M8BR and with Tyr-Tyr in MUT-16 FFR. Since Uebel et al.^29^ reported a loss of MUT-8 recruitment following the deletion of the M8BR region (633-772 aa), we further explored the recruitment of the MUT-8 N-terminal domain to the MUT-16 M8BR cluster using Martini3 and atomistic simulations. Martini3 simulations suggest that recruitment is primarily driven by interactions between the Tyr residues in the MUT-8 N-terminus and the positively charged Arg and Lys residues in MUT-16 M8BR. However, the atomistic simulations revealed a lower interaction propensity for Lys residues, further emphasizing the ‘stickiness’ of Lys residues observed in Martini3. The atomistic simulations highlighted the importance of cation-*π* or sp^2^-*π* interactions between Arg of MUT-16 M8BR and Tyr of MUT-8 N-terminal domain in the recruitment process. In particular, the atomistic simulations show that Arg-Tyr interactions are more stable than Lys-Tyr interactions, partly because Arg can form additional stabilizing hydrogen bonds with the Tyr backbone. *In vitro* pulldown experiments further support these findings, as mutating seven Arg residues in MUT-16 M8BR to Lys or Ala significantly reduced MUT-8 recruitment, confirming the key role of Arg residues in the recruitment of the MUT-8 N-terminal domain.

The ability to switch between different levels of resolution, from coarse-grained to atomistic models, will be increasingly crucial for studying large and complex biological systems. This approach captures the detailed chemical interactions and the broader molecular behaviours necessary for accurate simulations. In the case of *Mutator foci*, the same “molecular grammar”, a set of non-covalent interactions like cation-*π*,sp^2^-*π*, hydrogen bonds, plays a crucial role in regulating the behaviour of disordered proteins. These interactions, which have been well documented in the study of protein structure and dynamics, appear to govern the phase separation of MUT-16 and the recruitment of its client protein, MUT-8. By using multiscale simulations, we can better understand how these intricate molecular forces drive critical biological processes such as phase separation and protein recruitment, providing insights into the mechanisms that underpin cellular organisation and function.

## Supporting information

Supporting Information: The supplementary document provides additional data supporting the main findings, including Table S1, which details simulation

## Acknowledgement

This project was funded by SFB 1551 Project No. 464588647 of the DFG (Deutsche Forschungsgemeinschaft). L.S.S. acknowledges support by ReALity (Resilience, Adaptation and Longevity) and Forschungsinitiative des Landes Rheinland-Pfalz. A.C.S and L.S.S. thank M^3^ODEL for support. S.F. received funding from the Austrian Science Fund (FWF) programs I6110-B and the doc.funds DOC 177-B: RNA@core: “Molecular mechanisms in RNA biology”. V.B. received funding from the European Union’s Framework Programme for Research and Innovation Horizon 2020 (2014-2020) under the Marie Curie Skłodowska Grant Agreement Nr. 847548. (Vienna International PostDoc Program (VIP-2). We gratefully acknowledge the advisory services offered and the computing time granted on the supercomputers Mogon II at Johannes Gutenberg University Mainz, which is a member of the AHRP (Alliance for High-Performance Computing in Rhineland Palatinate) and the Gauss Alliance e.V. K.G. thanks Yashraj Wani, Rodrique Badr, and Lucia Baltz for insightful discussions. L.S.S. thanks, Dr. K. Lindorff-Larsen, Dr R. Sprangers, Dr. R. M. Bhaskara and Dr. J. Mittal, for inspiring discussions.

## Supporting Information Available

- MUT16_MUT8_interaction_Multiscale_Simulation_invitroexperiments_supplimentary.pdf: This PDF contains tables listing the simulations performed, disorder predictions, time-lapse of fusion of MUT-16 phase-separated droplets, region-wise contact maps, residuewise contact maps, density plots, and details of non-covalent interactions between Arg and Tyr.
- Mut16_FFR_phase_separation_timelapse.lif - Ce380_5-1_3.avi: This file contains a microscopy timelapse video of MUT-16 FFR droplet phase separation obtained *in vitro*.
- MUT16_M8BR_FFR_Martini3_simulation.mpg: This video showcases the simulation of spontaneous phase separation of MUT-16 M8BR+FFR chains.
- Atomistic_trajectory_of_MUT-16_and_MUT-8_interaction: The atomistic trajectory file detailing the interaction between MUT-16 and MUT-8, available on Zenodo (https://zenodo.org/uploads/12742133).

