## Supporting Information: The supplementary document provides additional data supporting the main findings, including Table S1, which details simulation for "Multi-scale Simulations of MUT-16 Scaffold Protein Phase Separation and Client Recognition"

Table S1: Simulation characteristics. Listed here are the various models and force-fields used to perform the simulations. Furthermore, the box dimensions, number of particles, type and number of proteins, and simulation time are also listed.

| Model | Force-field | Number of protein chains | Box dimension (nm) | Number of particles | Time |
|---|---|---|---|---|---|
| Atomistic | Amber99sb-star-ildn-q | 100 (MUT-16 M8BR)+10 (MUT-8) | 15×15×30 | 884932 | 1 µs |
| Atomistic (Folding@home) | Amber99sb-star-ildn-q | 100 (MUT-16 M8BR)+10 (MUT-8) | 15×15×30 | 884932 | 350 µs |
| Near-atomic coarse-grained | Martini3 (@$\lambda$ = 1,1.01,1.03,1.05) | 100 (MUT-16 M8BR+FFR) | 40×40×40 | 657674 | 10 µs |
| Near-atomic coarse-grained | Martini3 (@$\lambda$ = 1,1.01,1.03,1.05) | 180 (MUT-16 FFR) | 40×40×40 | 659058 | 10 µs |
| Near-atomic coarse-grained | Martini3 (@$\lambda$ = 1,1.01,1.03,1.05) | 220 (MUT-16 M8BR) | 40×40×40 | 658434 | 10 µs |
| Near-atomic coarse-grained | Martini3 | 100 (MUT-16 M8BR+FFR) | 60×60×60 | 1675095 | 20 µs |
| Near-atomic coarse-grained | Martini3 | 100 (MUT-16 M8BR+FFR)+20 (MUT-8) | 60×60×60 | 1676715 | 20 µs |
| Near-atomic coarse-grained | Martini3 | 100 (MUT-16 M8BR)+10 (MUT-8) | 15×15×60 | 138557 | 20 µs |
| Near-atomic coarse-grained | Martini3 (@$\lambda$ = 1.03) | 100 (MUT-16 M8BR)+10 (MUT-8) | 15×15×60 | 138557 | 20 µs |
| Residue-level coarse-grained | CALVADOS2 | 100 (MUT-16 M8BR +FFR) | 20×20×280 | 31200 | 10 µs |
| Residue-level coarse-grained | CALVADOS2 | 220 (MUT-16 M8BR) | 20×20×280 | 30800 | 10 µs |
| Residue-level coarse-grained | CALVADOS2 | 180 (MUT-16 FFR) | 20×20×280 | 30960 | 10 µs |
| Residue-level coarse-grained | CALVADOS2 (@$T = 260K, 265K, 270K, ..., 285K$) | 100 (MUT-16 M8BR +FFR) | 20×20×150 | 31200 | 10 µs |
| Residue-level coarse-grained | CALVADOS2 | 100 (MUT-16 M8BR +FFR)+10 (MUT-8) | 20×20×280 | 31710 | 10 µs |

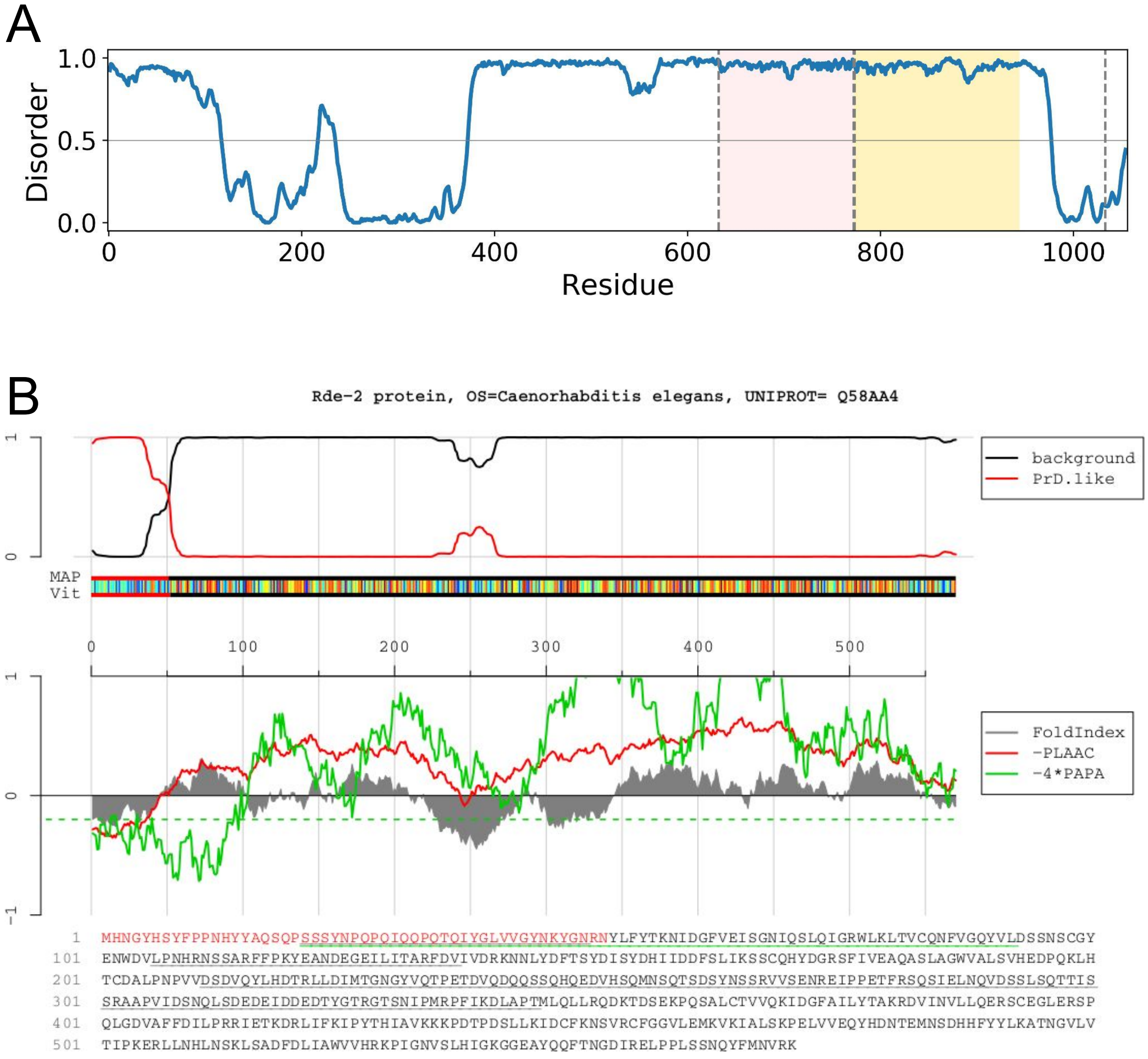


Figure S1: **A.** Disorder prediction for MUT-16, the M8BR (pink) and FFR (yellow) regions are highlighted. **B.** Prediction of the prion-like domain (PLD) in MUT-8 protein using PLAAC webserver[67]

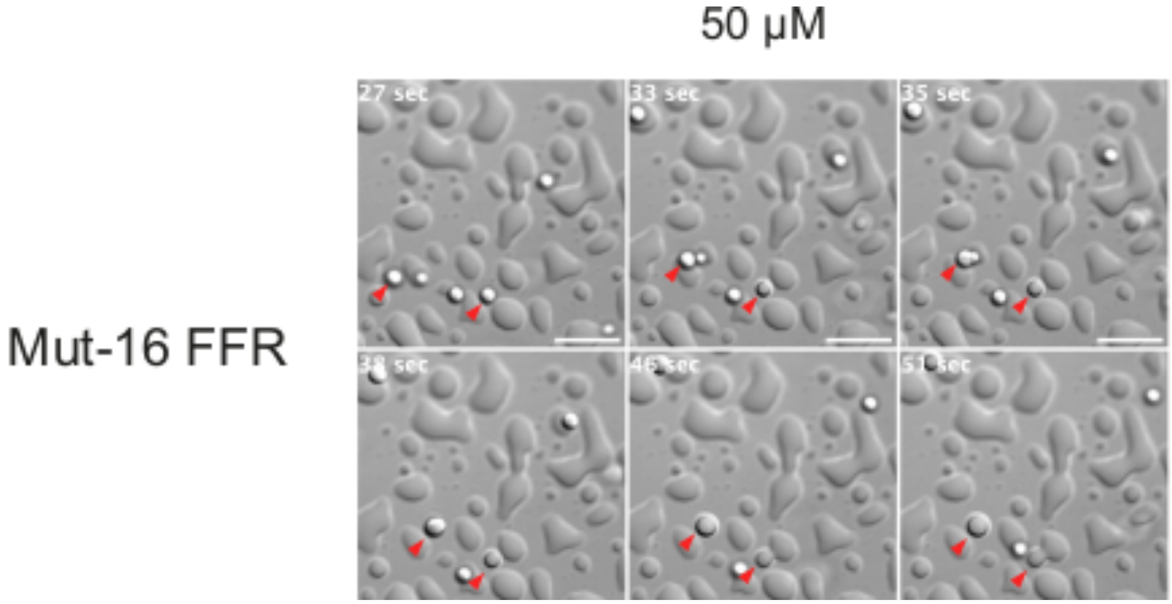


Figure S2: Time-lapse analysis demonstrates the fusion of phase-separated droplets formed by the addition of 50 µM MUT-16 FFR and a 1:100 ratio (w/w) of 3C protease to the MUT-16 fragment. Red arrowheads indicate droplets merging into larger ones and wetting the coverslip surface. Time refers to the imaging period. Scale bars: 10 µm.

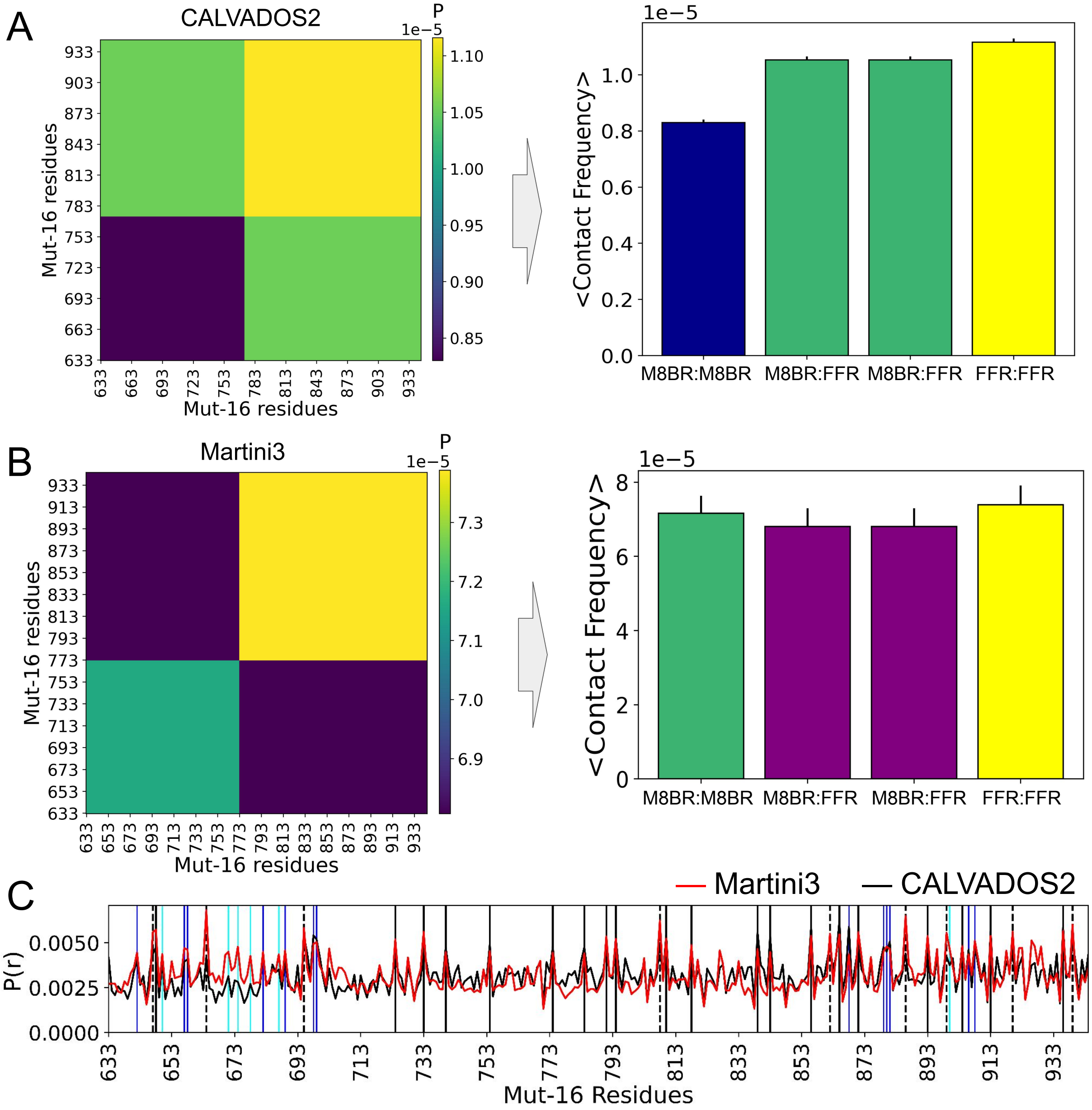


Figure S3: **A.** Region-wise contact frequency calculated for M8BR:M8BR (blue), M8BR:FFR (green) and FFR:FFR (yellow) interactions in residue-level coarse-grained simulation (CALVADOS2), Bar plot of average contact values from each region calculated from the CALVADOS2 simulations. **B.** Region-wise contact-map obtained for M8BR:M8BR (green), M8BR:FFR (purple) and FFR:FFR (yellow) interactions in near-atomic coarse-grained Martini3 simulations, Bar-plot of average contact frequency of interactions between different regions obtained through Martini3 simulations. **C.** One-dimensional contact map representing the peaks associated with the amino acid with higher relative contact probabilities (P(r)) obtained from CALVADOS2 and Martini3 simulations. The peaks shown by Tyr (black), Phe (dashed), Arg (blue), and Lys (cyan) are represented by the vertical lines.

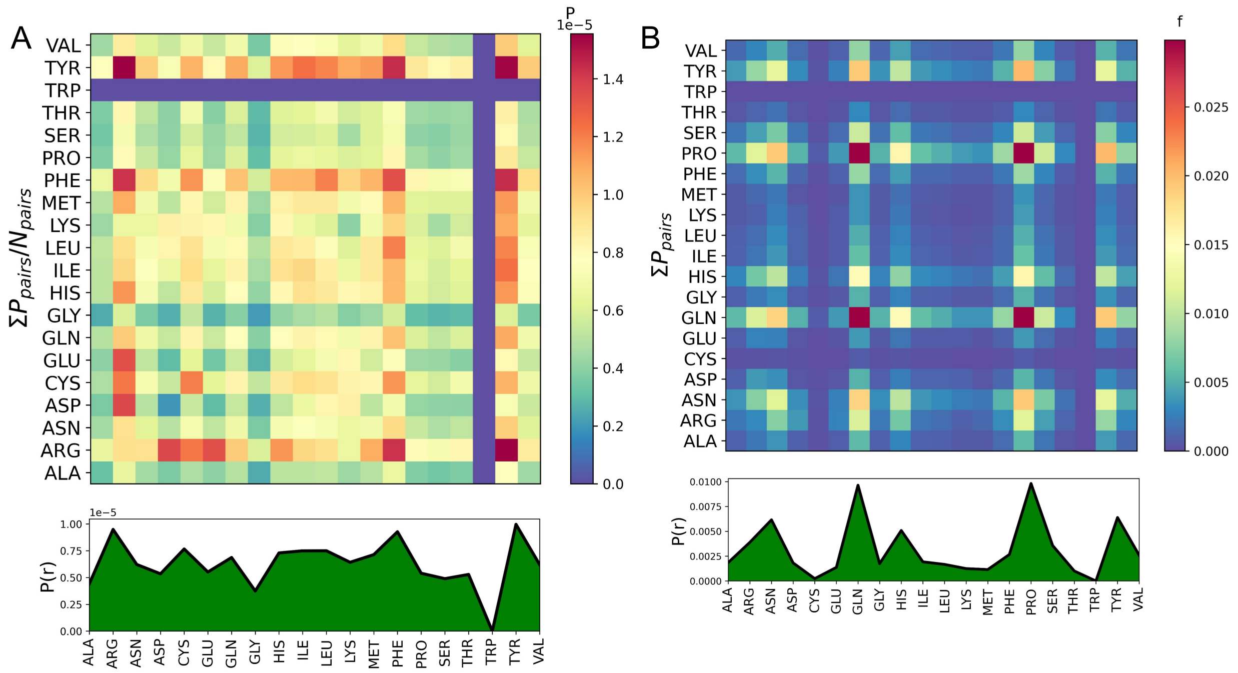


Figure S4: **A,B.** Normalised and Unnormalised amino-acid wise contact map of phase-separated MUT-16 M8BR+FFR chains obtained through the CALVADOS2 simulations at 275 K. Normalisation is done by the number of amino acid pairs in the system.

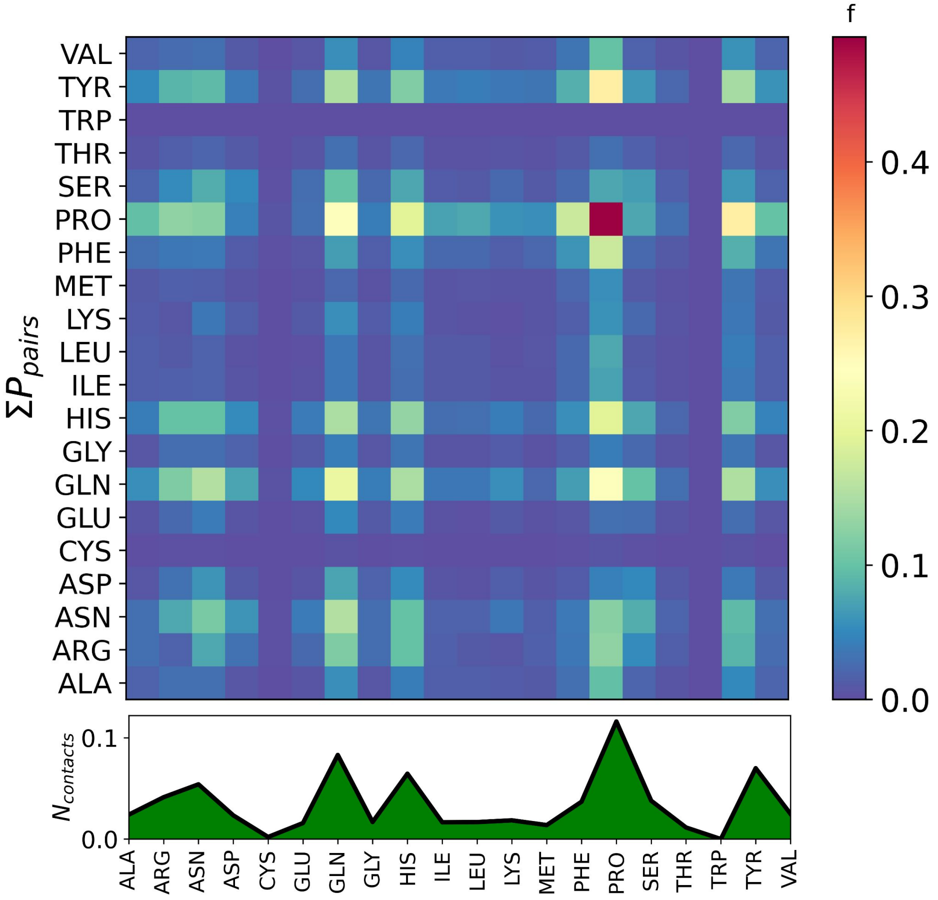


Figure S5: Unnormalised amino-acid wise contact map of phase-separated MUT-16 M8BR+FFR chains obtained through the Martini3 simulation.

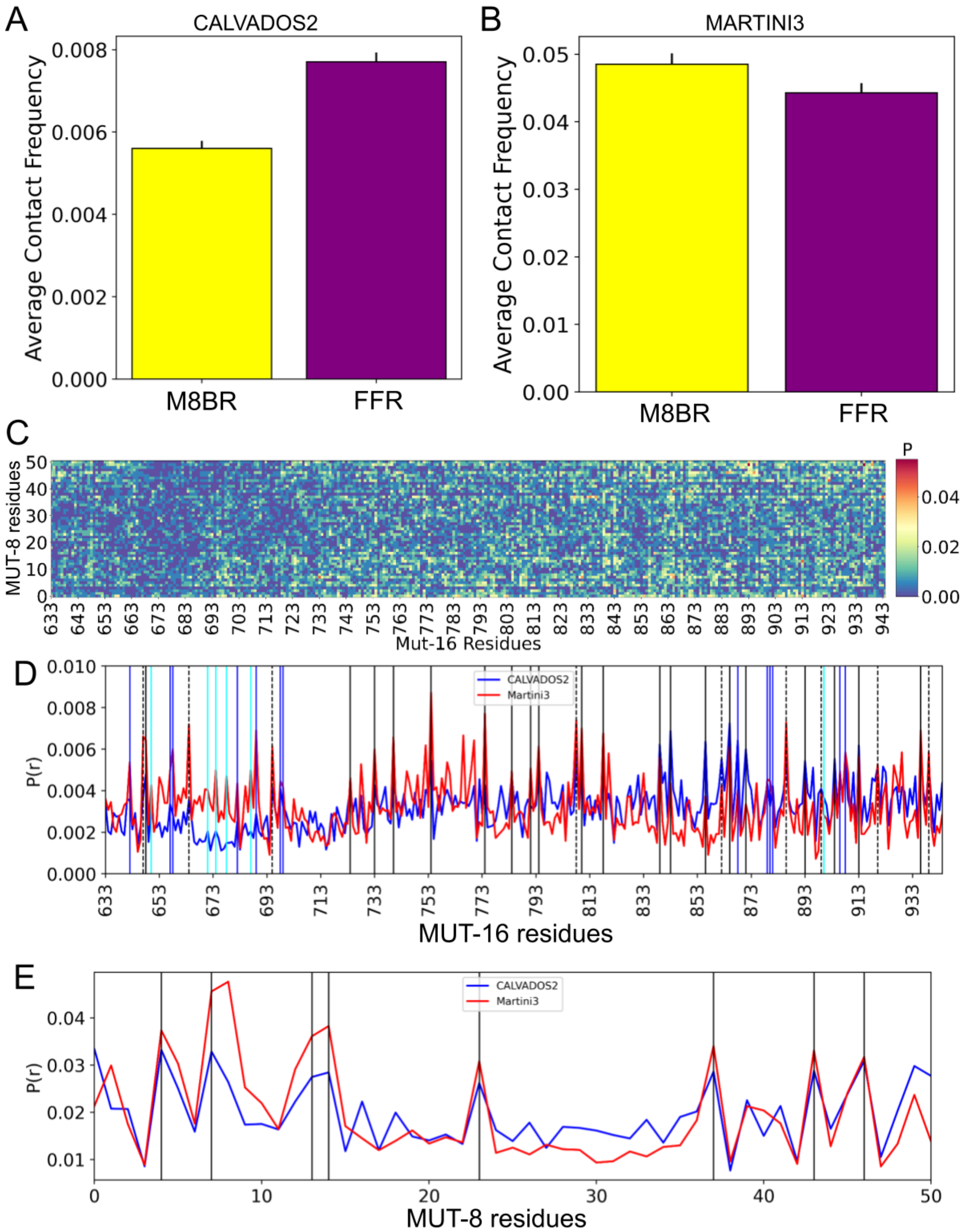


Figure S6: **A,B.** Average contact frequency of MUT-16 M8BR+FFR with MUT-8 N-terminal in CALVADOS2 and Martini3 simulations, respectively. **C.** 2-D contact map of interaction between MUT-8 and MUT-16 obtained from CALVADOS2 simulations. **D.** 1-D contact plot of MUT-16 (M8BR+FFR) obtained through Martini3 (red) and CALVADOS (blue), showing the relative frequencies of making contact per residue. Aromatic and positively charged amino acids like Tyr (black), Phe (black, dashed), Arg (blue), and Lys (cyan) are represented by the verticle lines. **E.** 1-D contact plot of MUT-8 N-terminal obtained through Martini3 (red) and CALVADOS2 (blue), showing the relative contact frequencies per residue.

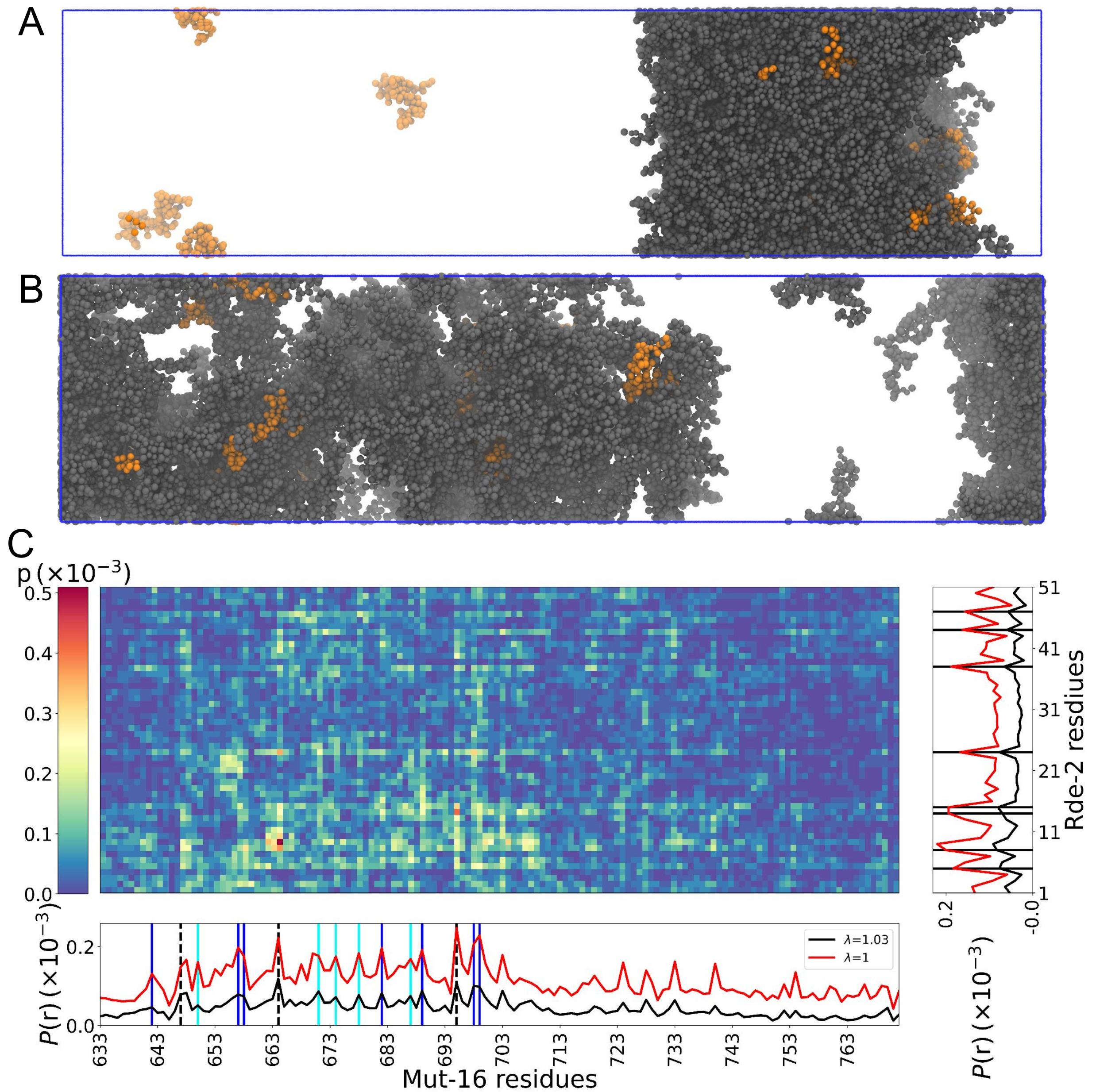


Figure S7: **A,B.**Final frame of the simulation for MUT-16 M8BR and MUT-8 N-terminal at $\lambda = 1$ and $\lambda = 1.03$. **C.** 2-D inter-molecular contact map of MUT-16 M8BR and MUT-8 at $\lambda = 1.03$. 1-D contact map at $\lambda = 1.03$ is compared to the simulation profile at $\lambda = 1$.

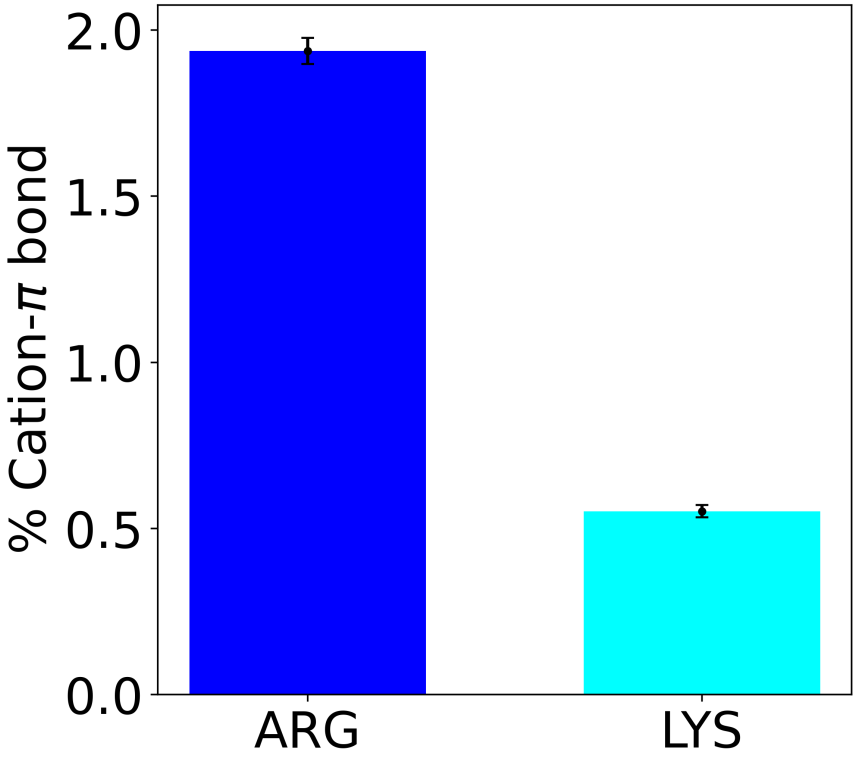


Figure S8: Percentage of cation-$\pi$ interactions between Arg/Lys of MUT-16 M8BR and Tyr of MUT-8 N-terminal domain with the Martini3 simulation model. Percentage is relative to the total amount of possible pairwise Arg and Lys Tyr interactions

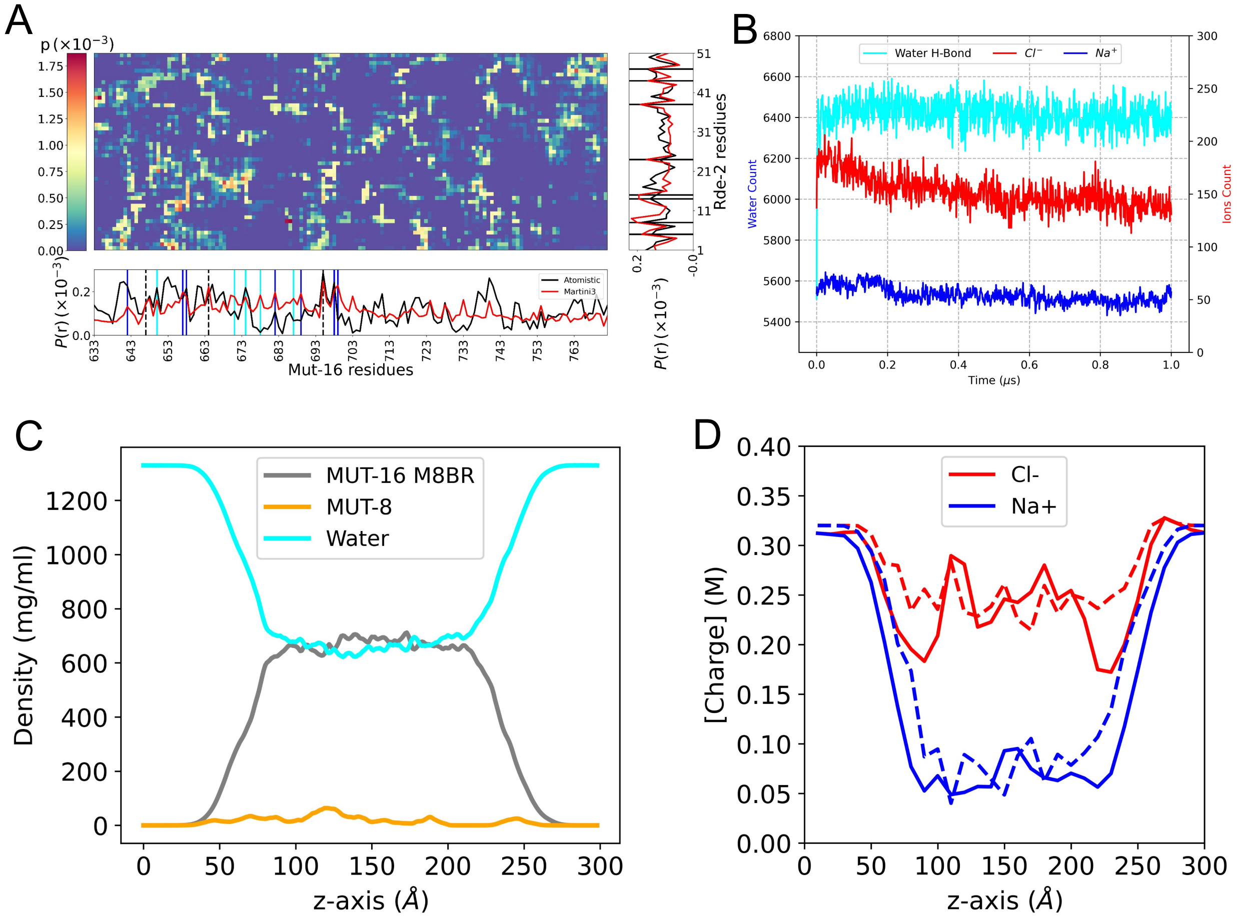


Figure S9: **A.** Contact map obtained from atomistic simulation of MUT-16 M8BR cluster and MUT-8 N-terminal PLD. 1-D contact map obtained by atomistic simulation and Martini3 simulations are compared. Residues like Tyr (black), Phe (dashed), Arg (blue), Lys (cyan) are indicated with vertical lines. **B.** Interactions of water and ions with MUT-16 M8BR during the 1 μs trajectory are shown. The cyan line represents the number of water molecules making hydrogen bonds with protein. The blue line represents the number of $Na^+$ ions bound to negatively charged residues. The red line represents the number of $Cl^-$ ions in electrostatic interaction with the positively charged residues. For calculating the number of ions bound an interaction a cutoff of 5 Å between the charged atoms was used. **C.** Density profiles obtained from atomistic simulations. Components are shown in the legend. **D.** Density profile corresponding to $Na^+$ and $Cl^-$ ions. Dashed lines represent the predicted values obtained from the concentration of cationic and anionic residues in both MUT-16 and MUT-8 proteins.

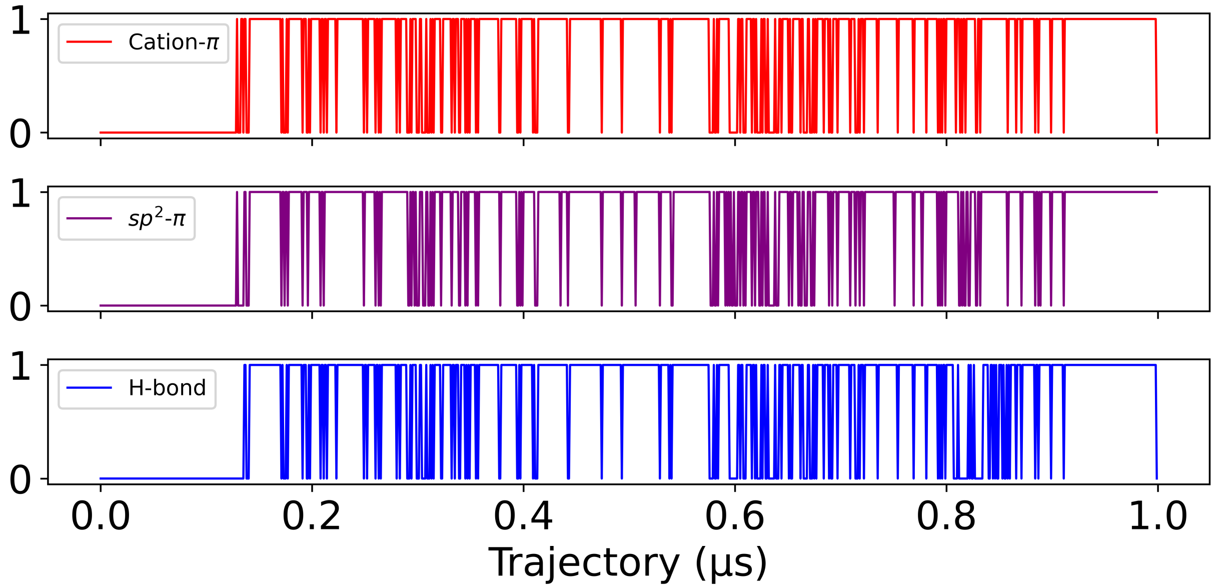


Figure S10: Different non-covalent interaction between a pair of Arg residues of MUT-16 M8BR and Tyr of MUT-8 N-terminal domain involving the cation-$\pi$, sp$^2$-$\pi$ and H-bond (given the cation-$\pi$ interaction has already formed).
